# EZH1 Q571R-mediated chromatin compaction and its oncogenic potential in thyroid cancer

**DOI:** 10.1101/2025.05.06.652476

**Authors:** Hanbyeol Kim, Do-Gyun Kim, Seongyun Ha, Jun-Young Jang, Francisca N. de L. Vitorino, Joanna M. Gongora, Woochul Kim, Yoonji Oh, Gyeongsik Park, Hyeji Kang, Soo-Hyun Im, Chan Kwon Jung, Seon-Young Kim, Inkyung Jung, Jae-kyung Won, Benjamin A. Garcia, Su-jin Kim, Kyu Eun Lee, Je-kyung Ryu, Chul-Hwan Lee

**Author notes:** These authors contributed equally as co-first authors (Hanbyeol Kim, Do-Gyun Kim, Seongyun Ha). Correspondence (CH.L.), prof.love@ snu.ac.kr (JK.R.), (KE.L).

## Abstract

Dysregulation of Polycomb Repressive Complex 2 (PRC2) is established to contribute to cancer. Of its catalytic subunits, EZH1 and EZH2, EZH2 mutations in cancer have been extensively identified and studied, but the role of EZH1 in cancer remains largely unexplored. The recurrent presence of the EZH1^Q571R^ mutation in follicular thyroid cancer suggests its involvement in tumor progression. Using EZH1 ChIP-seq, ATAC-seq, and H3K27me3 CUT&Tag, we demonstrated that EZH1^Q571R^ enhances chromatin binding and compaction and stimulates PRC2-EZH1 catalytic activity, leading to increased H3K27me3 levels and repression of tumor suppressor genes. Purified PRC2-EZH1^Q571R^ demonstrated a significant increase in histone methyltransferase activity compared to PRC2-EZH1^WT^ via enhanced nucleosome binding and DNA compaction. Notably, this effect is particularly observed with EZH1^Q571R^ but only to a lesser extent with the corresponding EZH2^Q570R^ mutation, highlighting it as an EZH1-specific mechanism. We additionally demonstrated PRC2-EZH1^Q571R^ to efficiently methylate H3K27 in pre-existing H3K36me2/3 nucleosomes, disrupting chromatin homeostasis. Our findings provide key insights into the molecular pathogenesis of EZH1^Q571R^-driven follicular thyroid cancer.

## Introduction

Chromatin regulation is essential in maintaining proper gene expression patterns throughout normal development; accordingly, its dysregulation could lead to developmental disorders and cancer. Among the various proteins involved in chromatin regulation, Polycomb Repressive Complex 2 (PRC2) is notable as a therapeutic target in multiple types of cancers.^1^ PRC2 plays a critical role in governing the formation of heterochromatin and gene silencing by catalyzing mono-, di-, and tri-methylation of histone H3 at lysine 27 (H3K27me1, -me2 and -me3).^2,3^ The complex consists of three core subunits, EED, SUZ12, and either of two mutually exclusive catalytic subunits, Enhancer of Zeste (EZH) 1 or 2.^2–4^ EZH1-containing PRC2 (PRC2-EZH1) and EZH2-containing PRC2 (PRC2-EZH2) exhibit functional differences in their catalysis of histone methylation and chromatin compaction.^5–9^ In particular, allosteric activation of PRC2-EZH2 is much efficient than that of PRC2-EZH1, whereas PRC2-EZH1 has higher nucleosome binding and chromatin compaction ability compared to PRC2-EZH2.

EZH2 mutations and overexpression have been linked to a variety of cancers, with high EZH2 expression frequently observed in solid tumors and associated with poor prognosis.^10–13^ Mutations at the Y646 position (Y646F, Y646N, Y646C, Y646S, and Y646H) are present in approximately 25% of diffuse large B-cell lymphoma cases, 12% of follicular lymphoma cases, and 1% of melanoma cases.^1^ Tazemetostat, a selective EZH2 inhibitor, has recently gained clinical approval for the treatment of follicular lymphoma and epithelioid sarcoma.^14^ However, several studies have shown that inhibiting EZH2 leads to increased expression of EZH1, which helps maintain H3K27 methylation levels and diminishes the therapeutic efficacy of EZH2 inhibition.^15,16^ Consequently, dual inhibition of both EZH1 and EZH2 has been explored, with the dual inhibitor Valemetostat currently undergoing clinical trials.^17^ However, where EZH2 has been extensively studied due to its well-established role in cancer cell proliferation, the relevance of EZH1 in cancer remains less well understood.

In the course of evolution, the ancestral Drosophila *E(z)* gene diversified into *EZH1* and *EZH2*.^18,19^ EZH2 shows high expression in early developmental stages, then significantly decreases during cellular differentiation.^20^ Its expression is regulated by E2F transcription factors, which are cell cycle-dependent, leading to the close association of EZH2 with cancer.^21^ Conversely, expression of EZH1 is low in actively dividing cells, including cancer cells,^1^ and correspondingly low in early development but greater in differentiated cells.^20^ Hence, EZH1 mutation has been only rarely found in cancer. Of particular note is the EZH1^Q571R^ mutation, in which a glutamine is replaced with an arginine at position 571, that was initially identified in about 27% of autonomous thyroid adenomas (ATAs).^22^ EZH1 mutations are the second most common genetic alteration in ATAs and are strongly associated with cAMP pathway genes, including activation of Thyroid-stimulating Hormone Receptor (TSHR).^22^ The EZH1^Q571R^ mutation was found to enhance the proliferation of normal thyroid cells, with more pronounced effect in the presence of an activating TSHR mutation.^22^ Since the initial discovery, EZH1 mutations have been increasingly observed in follicular thyroid adenomas and carcinomas.^22–25^ Another study discovered two hotspot mutations in thyroid cancers, EZH1^Q571R^ and EZH1^Y642F^, both of which were found to be driver mutations in follicular thyroid carcinoma and adenoma.^23^ Notably, EZH1^Y642F^ corresponds to EZH2^Y646F^, a well-known “kinetic” mutation found in various cancers.^26–28^ PRC2-EZH2^Y646F^ can convert H3K27me2 to H3K27me3 much more efficiently than the wild-type protein; potentially, EZH1^Y642F^ could have a similar effect. Meanwhile, the EZH1^Q571R^ mutation has no documented EZH2 counterpart, indicating a distinct molecular mechanism associated with this mutation in thyroid cancer. Recent studies have demonstrated that the EZH1^Q571R^ mutation results in elevated levels of H3K27me3, suggesting it to be a gain-of-function mutation.^22^ However, it is not well understood how this mutation could enhance the catalytic activity of PRC2.

In this study, we investigated the EZH1^Q571R^ mutant, utilizing a reconstituted system, isogenic thyroid epithelial cell lines, and thyroid cancer patient tissues to unveil the associated EZH1-specific cancer-driving mechanism. Our results demonstrate how EZH1^Q571R^ facilitates H3K27me3 invasion into euchromatic regions, leading to alterations in gene expression. These findings reveal the molecular patho-mechanisms of EZH1^Q571R^ in follicular thyroid cancer and offer new insights into the mutation’s oncogenic potential.

## Results

### Tumor tissue and thyroid epithelial follicular cell lines harboring the EZH1^Q571R^ mutation show elevation of H3K27me3

Building on the findings of Jung et al.^24^, which demonstrated that EZH1 mutations are predominantly found in oncocytic thyroid cancer (OC) and follicular thyroid cancer (FTC), we analyzed OC and FTC patient cohort data from Seoul National University Hospital (SNUH) and Catholic Medical Center (CMC) (Figures 1A and 1B). Among OC cases, EZH1^Q571R/K^ or EZH1^Y642F^ mutations were detected in 2 out of 5 patients from SNUH and 7 out of 29 from CMC (Figure 1A). Among FTC cases, these mutations were identified in 2 out of 69 patients from SNUH and 8 out of 77 from CMC (Figure 1B). Together, these results suggest strong associations of both mutations with both OC and FTC. Notably, in addition to the Q571R mutation, the Q571K mutation was also detected in OC. Since both of the substituted residues, arginine and lysine, carry a positive charge, they may enhance binding to negatively charged DNA (see below). We next sequenced 74 thyroid cancer patient tissues from the SNUH cohort and identified four cases (#1, #9, #52, and #53) heterozygous for the EZH1^Q571R^ mutation (Figures 1A, 1B, 1C and Table S1). While patients #1 and #9 had minimally invasive follicular thyroid carcinoma, patient #52 had oncocytic adenoma, and patient #53 oncocytic carcinoma. Immunohistochemistry analysis of the three carcinoma patients (#1, #9, and #53) revealed capsular invasion of cancer cells into surrounding normal tissue (Figure 1D), supporting the association of EZH1 mutations with thyroid cancer progression.

**Figure 1.**
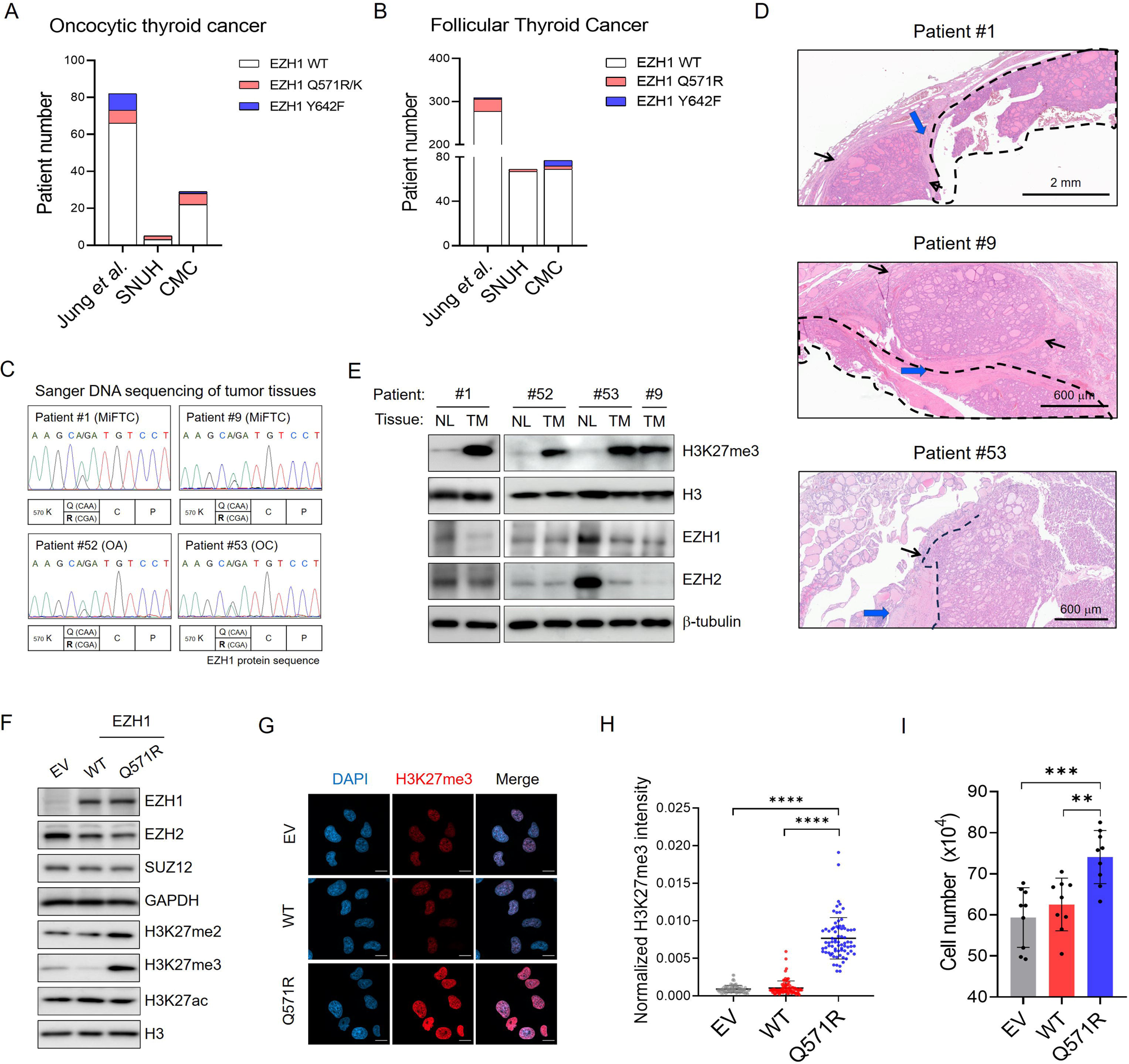
The EZH1^Q571R^ mutation, commonly found in thyroid cancer, enhances H3K27 methylation level. **(A-B)** Bar graph showing the number of oncocytic thyroid cancer patients (A) or follicular thyroid cancer patients (B) harboring EZH1 Q571R/K (red) or Y642F (blue) in Jung *et al.^24^*, SNUH, and CMC cohorts. **(C)** Sanger DNA sequencing confirming presence of the heterozygous EZH1^Q571R^ mutation in tumor tissues. **(D)** Immunohistochemistry images of thyroid cancer tissues harboring EZH1^Q571R^ mutation. Dashed black lines indicate the boundaries of the tumor region. Blue arrows indicate fibrous capsule and black arrows indicate capsular invasion of cancer cells. Scale bar: 2 mm (top), 600 µm (middle), and 600 µm (bottom). **(E)** Western blot of H3K27me3, H3, EZH1, EZH2, and β-tubulin in matched normal (NL) and thyroid tumor (TM) tissues harboring EZH1^Q571R^ mutation. **(F)** Western blot of EZH1, EZH2, SUZ12, GAPDH, H3K27me2, H3K27me3, H3K27ac, and H3 in thyroid follicular epithelial cells (Nthy-ori-3-1) constitutively expressing empty vector (EV), EZH1^WT^, or EZH1^Q571R^. **(G)** Representative immunofluorescence images of Nthy cells expressing EV (top), EZH1^WT^ (middle), or EZH1^Q571R^ (bottom). Cells were stained with DAPI and H3K27me3 antibody. Scale bar: 20 µm. **(H)** Quantification of H3K27me3 intensity normalized to nuclear area. Data are presented as mean ± SEM for EV (n=108), EZH1^WT^ (n=99), and EZH1^Q571R^ (n=96). Statistical differences were assessed using two-tailed Welch’s *t*-test (\*\*\*\**p*<0.0001). **(I)** Bar graph depicting the number of Nthy cells expressing EV, EZH1^WT^, or EZH1^Q571R^. 20,000 cells were seeded into 10-cm plates, and counts were taken after 48 hours. Data are shown as mean ± SEM from three independent experiments. Statistical differences were assessed using two-tailed Welch’s *t*-test (\*\**p*<0.01 and \*\*\**p*<0.001).

To explore the influence of the EZH1^Q571R^ mutation on H3K27 methylation in tumor tissue, we analyzed paired tumor and normal tissue samples from three patients (#1, #52, and #53). For patient #9, only tumor tissue was available (Figure 1E). Western blotting was performed to assess protein expression and H3K27 methylation. Although EZH1 and EZH2 expression levels in tumor tissue remained unchanged or reduced compared to normal tissue, tumor tissue carrying the EZH1^Q571R^ mutation exhibited a significant increase in H3K27me3 level (Figure 1E). This suggests that the EZH1^Q571R^ mutation significantly alters PRC2 complex activity, leading to enhanced H3K27 trimethylation.

To confirm the increased histone methyltransferase activity of PRC2-EZH1^Q571R^ in an isogenic condition, we generated cell lines in ‘Nthy-ori-3-1’, a human thyroid follicular epithelial cell line, with constitutive expression of either empty vector, EZH1^WT^, or EZH1^Q571R^. The overexpression of EZH1^Q571R^ resulted in a significant increase in global H3K27me3 level, evident in both western blot and immunofluorescence imaging (Figures 1F-1H and S1A), suggesting the EZH1^Q571R^ mutation to significantly increases the histone methyltransferase (HMT) activity of PRC2. Notably, mass spectrometry analysis showed that this mutation did not alter levels of other repressive histone modifications such as H3K9me3 and H4K20me3 (Figure S1B).

To evaluate whether the EZH1^Q571R^ mutation promotes cell proliferation, we examined and tracked cell growth. Nthy-ori-3-1 cells with induced expression of EZH1^Q571R^ exhibited enhanced proliferation compared to cells with induced expression of EZH1^WT^ (Figure 1I), indicating the proliferative role of this mutation.

### Cells expressing the EZH1^Q571R^ mutant display increased chromatin condensation, elevated H3K27me3, and repression of tumor suppressor genes

To investigate how EZH1^Q571R^ modifies chromatin structure and epigenetic states, we performed EZH1 ChIP-seq, H3K27me3 CUT&Tag, and ATAC-seq in Nthy cell lines expressing either EZH1^WT^ or EZH1^Q571R^ (Figures 2A and S2A-S2C). Compared to EZH1^WT^, EZH1^Q571R^ resulted in greater chromatin occupancy (Figure 2A). Consequently, regions where EZH1^Q571R^ is enriched exhibited significant increases in H3K27me3 level and compacted chromatin (Figure 2A). Collectively, these findings suggest that greater binding of EZH1^Q571R^ to chromatin enhances H3K27 methylation and promotes chromatin compaction.

**Figure 2.**
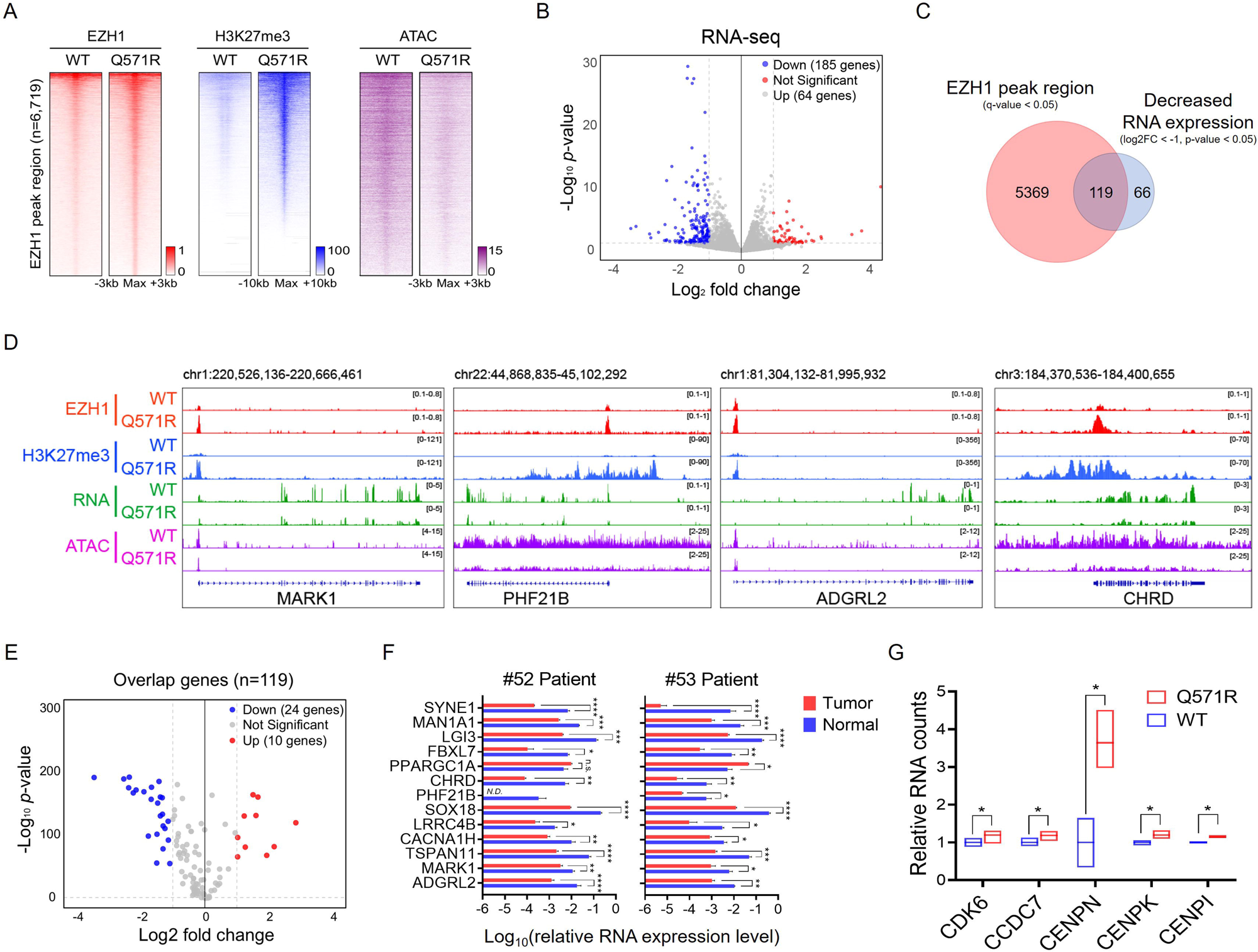
Cells with EZH1^Q571R^ show stronger chromatin binding, increased H3K27me3, chromatin condensation, and tumor suppressor gene suppression. **(A)** Heatmaps displaying EZH1 ChIP-seq, H3K27me3 CUT&Tag, and ATAC-seq (chromatin accessibility) signals across 6,719 EZH1 peak regions in Nthy cell lines expressing either EZH1^WT^ or EZH1^Q571R^. Each row represents a 4 kb window for EZH1, a 10 kb window for H3K27me3, or a 6 kb window for ATAC-seq, centered on the peak’s maximum signal. The signal intensity represents the average value from two replicates. **(B)** Volcano plot illustrating differential gene expression in Nthy cell lines expressing EZH1^WT^ or EZH1^Q571R^. A total of 185 genes (blue) were significantly downregulated, while 64 genes (red) were significantly upregulated in Nthy cells expressing EZH1^Q571R^ compared to EZH1^WT^. Genes without significant change are shown in gray. The x-axis represents the log_2_ fold change, and the y-axis represents the −log_10_ *p*-value; the significance threshold was set at log_2_|Fold Change| > 1 and −log_10_P.value < 1.301. Three replicates of each sample were used in the analysis. **(C)** Venn diagram showing overlap of the 5,488 genes near EZH1 peak regions in EZH1^Q571R^-expressing Nthy cell lines (red circle) and the 185 genes significantly downregulated in the same cell lines (blue circle). The overlap of the two sets comprised 119 genes. **(D)** Representative Integrative Genomics Viewer tracks illustrating EZH1 (red), H3K27me3 (blue), RNA-seq (green), and chromatin accessibility (purple) signals in Nthy cell lines expressing EZH1^WT^ or EZH1^Q571R^. The selected genes are known tumor suppressors downregulated in the EZH1^Q571R^ mutant condition. **(E)** Volcano plot of the 119 overlap genes from Panel C illustrating expression differences between 513 thyroid tumor tissues (TCGA) and 686 thyroid normal tissues (GTEx). A total of 24 genes (blue) were significantly downregulated in tumor tissues, while 10 genes (red) were significantly upregulated. Genes without significant changes are shown in gray. The x-axis represents log_2_ fold change, and the y-axis represents −log_10_ *p*-value; the significance threshold was set at log_2_|Fold Change| > 1 and −log_10_P.value < 2. **(F)** Bar graph illustrating the RT-qPCR results of 13 EZH1^Q571R^-downregulated tumor suppressor genes in tissues from patients #52 and #53. Data are presented as mean ± SEM, n=3 independent experiments. Statistical differences were assessed using two-tailed Welch’s *t*-test (\**p*<0.05, \*\**p*<0.01, \*\*\**p*<0.001, \*\*\*\**p*<0.0001, and ns: not significant). **(G)** Box plot representing relative normalized RNA counts of genes known to promote cell proliferation, which are upregulated in EZH1^Q571R^-expressing Nthy cells. Data are presented as mean ± SEM, n=3 independent experiments. Statistical differences were assessed using DESeq2 (\**p*<0.05).

To identify genes silenced due to increased H3K27me3 and chromatin condensation in Nthy cells expressing EZH1^Q571R^, we performed RNA sequencing (Figure S2D). In EZH1^Q571R^-expressing cells compared to wild-type EZH1^WT^, our analysis revealed that 64 genes were significantly upregulated, while 185 were downregulated (Figure 2B). Among those downregulated genes, 119 (64%) were directly influenced by EZH1^Q571R^ (Figures 2C and 2D).

Next, we investigated the expression of these directly-influenced genes in both normal and thyroid tumor tissues by analyzing gene expression data from GTEx (686 normal thyroid samples) and TCGA (513 thyroid tumor samples). Among the 119 genes, 24 were significantly downregulated in thyroid tumors (Figures 2E and S2E). Notably, 13 of these genes are recognized tumor suppressors, and their downregulation was further confirmed in tumor tissues from patients #52 and #53, with the exception of *PPARGC1A* (Figure 2F).

The RNA sequencing also revealed 5,369 genomic regions bound by EZH1 that did not show significant changes in RNA expression. This lack of response was attributed to the naturally low transcription levels of these regions under normal physiological conditions (Figure S2F). However, KEGG analysis indicated genes within these regions to be primarily associated with cancer-related pathways, including ‘transcriptional misregulation in cancer’ and ‘proteoglycans in cancer’ (Figure S2G), with enriched pathways including key tumor suppressor genes such as *WT1*, *WIF1*, and *HPGD* that remained suppressed in EZH1^Q571R^-expressing cells due to persistent H3K27me3 accumulation (Figure S2H). Finally, although this effect may be indirect, several genes associated with cell proliferation were significantly upregulated in EZH1^Q571R^-expressing Nthy cells, suggesting a shift toward a proliferative transcriptional program (Figure 2G). Collectively, these findings indicate that EZH1^Q571R^ may promote cell proliferation by repressing the expression of tumor suppressor genes and indirectly activating cell cycle-related genes.

### PRC2-EZH1^Q571R^ does not alter intrinsic catalytic activity but significantly increases nucleosome binding and H3K27me3

Recent cryo-EM studies have revealed important insights into the positioning of the EZH1 Q571 residue within the dimeric PRC2-EZH1 complex in relation to nucleosomal DNA.^8^ Specifically, EZH1 Q571 is situated within the major groove of the nucleosomal DNA (Figures 3A and 3B), and the two Q571 residues of the dimer have the capacity to interact with the DNA at the nucleosome exit site (Figure 3B). This interaction may serve to stabilize the nucleosome structure and promote efficient chromatin compaction.

**Figure 3.**
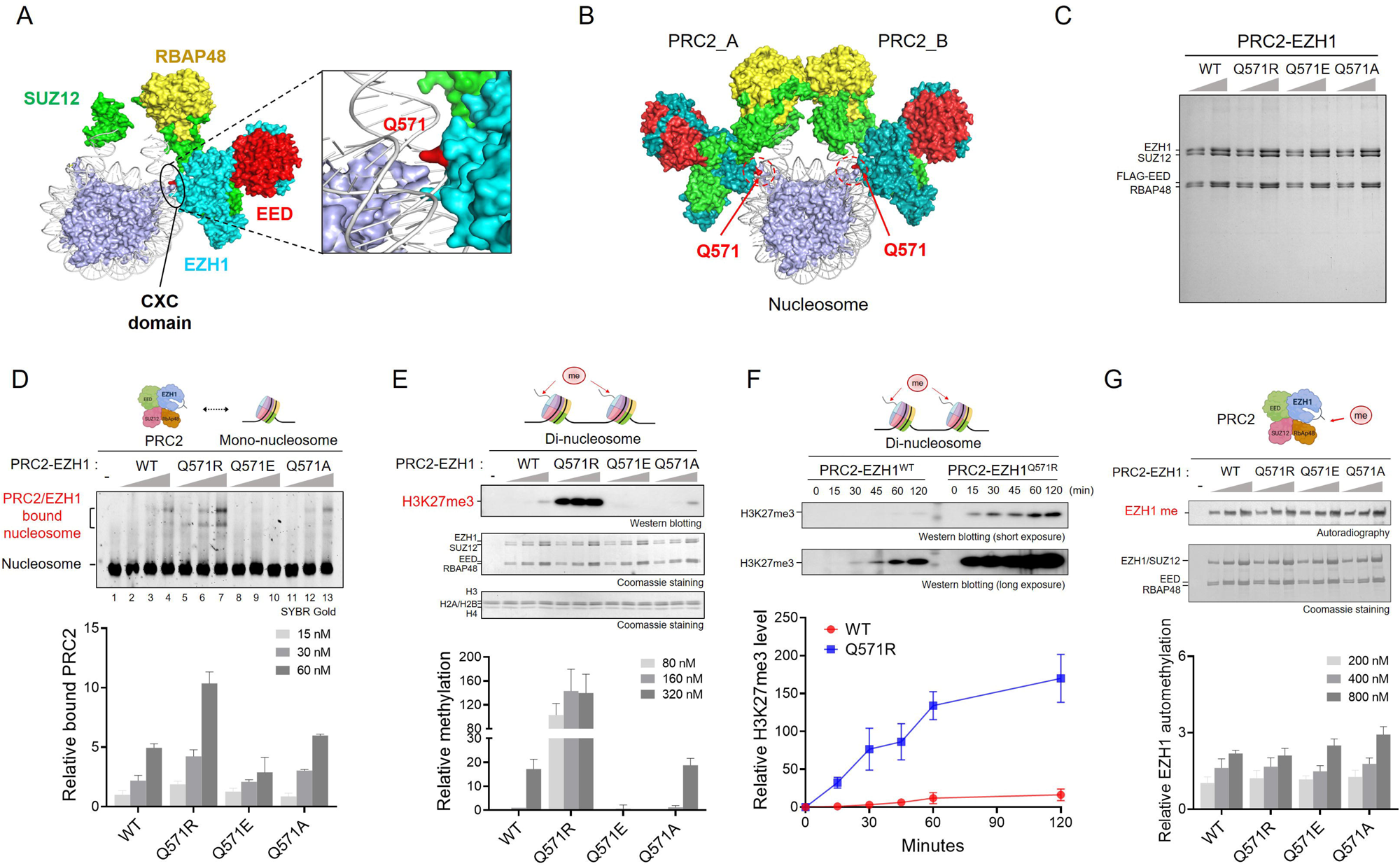
PRC2-EZH1^Q571R^ significantly stimulates histone methyltransferase activity via enhanced nucleosome binding. **(A)** Left: Cryo-EM structure showing the PRC2-EZH1 complex bound to a mono-nucleosome (modified from mono-nucleosome [PDB: 7KTQ] and PRC2-EZH1 complex [PDB: 7KTP]). The black oval highlights the CXC domain of EZH1. Right: Close-up showing the CXC domain of EZH1. The residue colored red corresponds to EZH1 Q571. **(B)** Cryo-EM structure of the dimeric form of PRC2-EZH1bound to a mono-nucleosome (modified from mono-nucleosome [PDB: 7KTQ] and PRC2-EZH1 complexes [PDB: 7KTP and 7KSR]). Red residues inside dashed circles correspond to EZH1 Q571 for each PRC2 complex. **(C)** Coomassie blue staining of SDS-PAGE gels showing purified recombinant PRC2 subunits (SUZ12, EED, RBAP48) along with WT or mutant EZH1 (Q571R, Q571E, or Q571A). **(D)** Electrophoretic mobility shift assay (EMSA) results. Data are presented as mean ± SEM, n=3 independent experiments. Top: Representative EMSA gel showing the binding affinity of PRC2 complex (15, 30, or 60 nM) to 80 ng mono-nucleosomes. Bottom: Quantification of the relative level of PRC2 complex bound to mono-nucleosomes. **(E)** Histone methyltransferase (HMT) assays. Data are presented as mean ± SEM, n=3 independent experiments. Top: HMT assays were conducted in the presence of 300 ng di-nucleosome and varying concentrations of PRC2 (80, 160, or 320 nM); H3K27me3 level was assessed by western blot (top image), and the relative amounts of PRC2 components (middle image) and di-nucleosomes (bottom image) by Coomassie blue staining of SDS-PAGE gels. Bottom: Quantification of the relative level of H3K27me3-modified di-nucleosomes. **(F)** H3K27me3 modification over time (0, 15, 30, 45, 60, and 120 minutes) as determined by HMT assays using 300 ng of di-nucleosome and 400 nM of PRC2. Data are presented as mean ± SEM, n=3 independent experiments. Top: Representative western blot measuring H3K27me3 level. Bottom: Quantification of the relative level of H3K27me3-modified di-nucleosomes. **(G)** Determination of EZH1 automethylation level. Data are presented as mean ± SEM, n=3 independent experiments. Top: HMT assays were conducted with varying concentrations of PRC2 (80, 160, or 320 nM); incorporation of [^3^H]-SAM into EZH1 was quantified by autoradiography (top image), and the relative amount of PRC2 components by Coomassie blue staining of SDS-PAGE gels (bottom image). Bottom: Quantification of the relative level of [^3^H]-SAM-incorporated EZH1.

Based on this observation, we hypothesize that the EZH1^Q571R^ mutation enhances the interaction with negatively charged nucleosomal DNA due to the positive charge of the arginine residue. This strengthened interaction may facilitate chromatin compaction, leading to increased H3K27 methylation.

To test this hypothesis, we performed an electromobility shift assay assessing the nucleosome binding activity of purified PRC2-EZH1^WT^ and mutant complexes: PRC2-EZH1^Q571R^, PRC2-EZH1^Q571E^, and PRC2-EZH1^Q571A^ (Figure 3C). We classified the Q571E mutation as introducing a negative charge, in contrast to the positively charged Q571R, while Q571A was considered a neutral substitution. Notably, PRC2-EZH1^Q571R^ exhibited stronger nucleosome binding than PRC2-EZH1^WT^, whereas PRC2-EZH1^Q571E^ displayed a significant defect in nucleosome binding (Figure 3D). These findings indicate that the presence of a positively charged arginine at this position is crucial for enhanced nucleosome binding. Interestingly, a Q571K mutation has also been identified in thyroid cancer patients (Figure 1A), suggesting that substitution of glutamine (Q) with a positively charged residue (R or K) at this position is associated with increased PRC2 activity and may contribute to thyroid cancer development.

Next, we conducted an *in vitro* histone methyltransferase (HMT) assay to evaluate the catalytic activity of PRC2-EZH1. Our results revealed a significant increase of HMT activity in the PRC2-EZH1^Q571R^ mutant, whereas the PRC2-EZH1^Q571E^ mutant exhibited reduced activity compared to the WT complex (Figures 3E, 3F and S3A), congruent with the nucleosome binding results. When using an octamer lacking nucleosomal DNA as a substrate, the enhancement observed with PRC2-EZH1^Q571R^ was abolished (Figure S3B), supporting that the interaction between EZH1 Q571R and nucleosomal DNA is necessary for the enhanced catalytic activity. Finally, to determine whether the EZH1^Q571R^ mutation directly enhances the intrinsic activity of PRC2, we examined EZH1 automethylation levels.^29,30^ Interestingly, the mutation had no effect on EZH1 automethylation (Figure 3G). Taken together, these results suggest that the PRC2-EZH1^Q571R^ mutant selectively enhances histone methyltransferase activity by strengthening nucleosome binding without altering intrinsic catalytic activity.

### PRC2-EZH1^Q571R^ exhibits enhanced DNA compaction compared to the wild-type

Although the PRC2-EZH1^Q571R^ mutant complex exhibits both increased nucleosome binding and enhanced HMT activity, the degree to which PRC2 activity is stimulated (>15-fold) far exceeds the modest twofold increase in nucleosome binding (Figures 3D and 3E). This difference led us to hypothesize that EZH1^Q571R^ might promote DNA compaction, creating a more condensed chromatin environment by reducing the length of DNA linker. Such dense chromatin could serve as a more favorable substrate for H3K27 methylation, as suggested by previous studies.^31,32^ To test this, we conducted a DNA compaction experiment using a single-molecule fluorescence assay^33^ in which we applied PRC2 onto λ-DNA immobilized on a PEG-coated surface and observed the formation of clustered spots (Figures 4A). To quantify the degree of DNA compaction, we compared consecutive image frames for fluctuation in DNA clustering (Figures S4A and S4B, and Methods and Materials). While the DNA alone exhibited constant fluctuation, addition of PRC2 led to an exponential decrease in fluctuation (Figures 4B and 4C). Remarkably, we observed the PRC2-EZH1^Q571R^ mutant to compact DNA at a faster rate compared to the wild-type, whereas the PRC2-EZH1^Q571E^ mutant showed no binding and no compaction (Figures 4D and S4C-S4G). These results support that the stimulating effect of EZH1^Q571R^ is due to an electrostatic interaction between the positively charged arginine residue and negatively charged DNA.

To further demonstrate that PRC2-EZH1^Q571R^ facilitates and sustains DNA compaction more effectively than the WT and other EZH1 mutants, we performed a magnetic tweezer experiment^34,35^ using a 9.2 kbp fragment of double-stranded DNA conjugated with a magnetic bead and a glass surface (Figure 4E). At a constant force of 3 pN, the DNA remained fully extended. PRC2 was introduced, and after 50 seconds, the force was reduced to 0.15 pN, leading to DNA compaction. This compaction was observed as a reduction in the fragment’s end-to-end length, driven by PRC2-mediated condensation. We quantified the degree of DNA compaction by measuring Z-bead positions under control conditions (without protein) and in the presence of PRC2. At 0.15 pN, the bead position reached approximately −1 µm in the absence of protein. However, in the presence of PRC2, it shifted to approximately −1.7 µm (Figure 4F), indicating DNA compaction.

**Figure 4.**
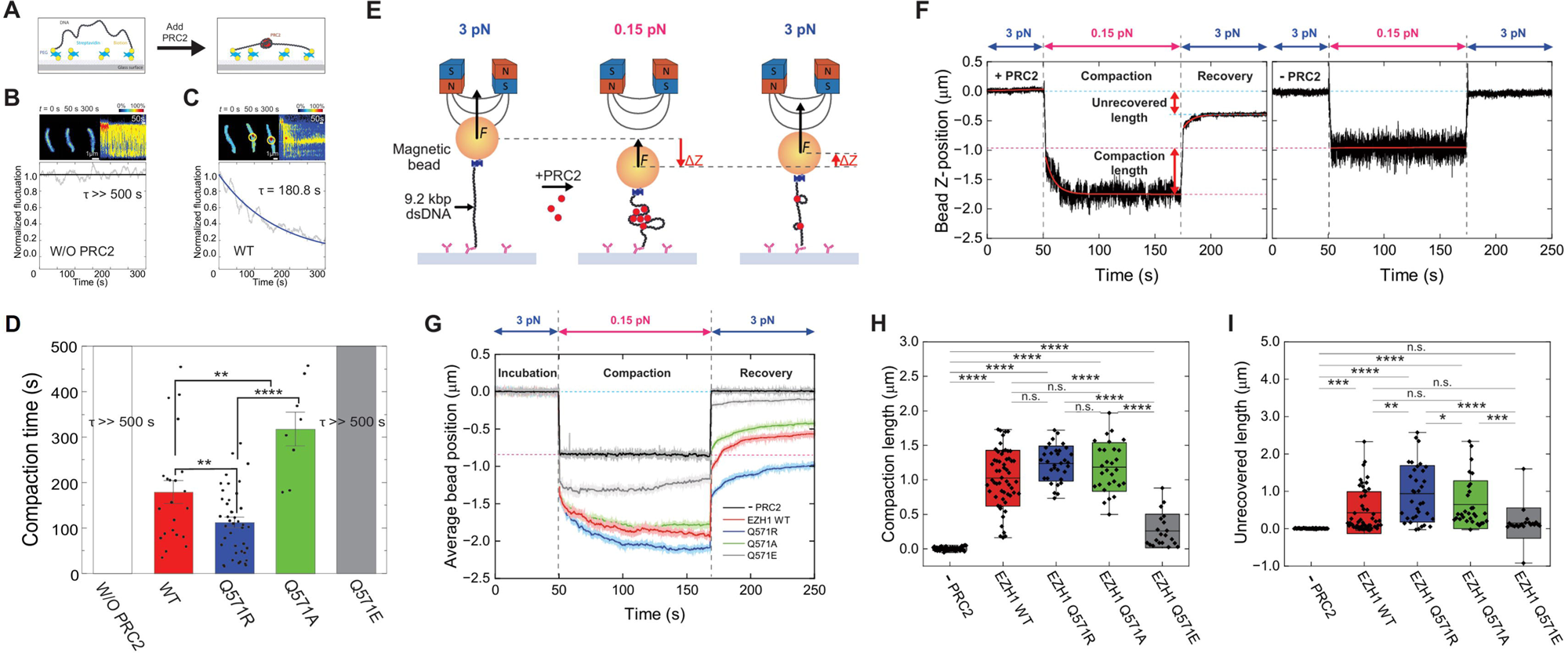
PRC2-EZH1^Q571R^ compacts DNA much more efficiently than the wild-type. **(A)** Schematic depicting single-molecule imaging of the DNA tether assay to observe DNA compaction by PRC2, before (left) and after (right) DNA compaction by PRC2. **(B-C)** Real-time DNA compaction by PRC2: (B) Control data without PRC2. (C) DNA compaction after PRC2 arrival. Top-left images are snapshots and the top-right the kymograph. The red star indicates when compaction starts. Bottom graphs show the fluctuation of DNA. Solid blue lines indicate a single-exponential fit with a time constant, τ. **(D)** Boxplot showing compaction time of PRC2-EZH1^WT^, -EZH1^Q571R^, -EZH1^Q571A^, and -EZH1^Q571E^ (mean ±SEM, *n* = 30, 22, 42, 8, 6). Significant difference determined by two-tailed Student’s *t*-test (\*\**p*<0.01 and \*\*\*\**p*<0.0001). **(E)** Experimental scheme depicting the magnetic tweezers experiment. PRC2 was applied under an initial high force (3 pN), which was decreased to 0.15 pN to induce DNA compaction. Then, higher force (3 pN) was reapplied to reverse compaction. **(F)** Representative single-molecule trajectory of DNA compaction and recovery with PRC2 (left) and without PRC2 (right). Black lines represent the raw data, red lines show the data fitted to an exponential function for the low force region, cyan dotted lines denote positions before DNA compaction, and red dotted lines indicate positions during compaction. **(G)** Average trace of DNA compaction and recovery in the absence of PRC2 or in its presence with EZH1 WT, Q571R, Q571A, or Q571E. Thick lines at full color opacity represent filtered graphs using a moving median filter (0.9 Hz), while lower-opacity halos represent the raw data. **(H-I)** Plots of DNA compaction length (H) and unrecovered DNA length (I), defined in (F). N = 57 for no PRC2 and 57, 35, 31, and 22 for PRC2 possessing EZH1 WT, Q571R, Q571A, and Q571E, respectively. Box plots represent the range from mean – s.d. to mean + s.d.; center lines show means and whiskers the minimum and maximum values. *P*-values were assessed using a two-tailed Student’s *t*-test (\**p*<0.05, \*\**p*<0.01, \*\*\**p*<0.001, \*\*\*\**p*<0.0001, and ns: not significant).

Further comparison among the different complexes revealed DNA to be compacted more effectively by PRC2-EZH1^Q571R^ than by the WT or other mutant complexes (Figure 4G), consistent with the results of single-molecule fluorescence assays. Once compaction began, a gradual exponential decrease in bead position was observed, achieving DNA compaction lengths (mean ± s.d.) of 1.02 ± 0.40 µm for PRC2-EZH1^WT^, 1.24 ± 0.25 µm for PRC2-EZH1^Q571R^, 1.18 ± 0.35 µm for PRC2-EZH1^Q571A^, and 0.26 ± 0.24 µm for PRC2-EZH1^Q571E^ (Figure 4H). Notably, where PRC2-EZH1^Q571R^ was the most effective, the negatively charged PRC2-EZH1^Q571E^ mutant showed minimal compaction, suggesting that electrostatic interactions between PRC2 and DNA play a crucial role (Figure 4H).

Maintenance of DNA compaction was examined under high DNA-stretching force (3 pN). The unrecovered DNA length (the difference between the final extended position and the original full-length position) was negligible in the PRC2-free control but varied in PRC2-induced compacted DNA according to EZH1 mutant status, with PRC2-EZH1^WT^, -EZH1^Q571R^, -EZH1^Q571A^, and - EZH1^Q571E^ achieving lengths of 0.43 ± 0.56, 0.93 ± 0.76, 0.64 ± 0.64, and 0.00 ± 0.01 µm, respectively (Figure 4I). Whereas the PRC2-EZH1^Q571E^ mutant rapidly reverted to its original extended state, PRC2-EZH1^Q571R^ retained the most compacted DNA structure even under high force, suggesting that the electrostatic interaction between EZH1^Q571R^ and DNA is exceptionally strong and resistant to disruption (Figure 4I).

### PRC2-EZH1^Q571R^ alters actively transcribed regions marked by H3K36me2/3

We next investigated whether the PRC2-EZH1^Q571R^ mutant could overcome the inhibition of H3K27 methylation in the presence of active histone modifications, particularly H3K36me2 and H3K36me3. PRC2 activity is known to be suppressed when H3K36me2/3 is present on the same histone,^36,37^ a well-documented antagonistic relationship that plays a crucial role in maintaining proper gene expression patterns by ensuring the segregation of euchromatin and heterochromatin. A recent cryo-EM structural study showed that H3K36me3 and the EZH2^Q570^ residue are positioned within the same groove of nucleosomal DNA, suggesting that H3K36me3 inhibits PRC2 by direct competition.^38^ Based on this, we hypothesized that PRC2-EZH1^Q571R^ might overcome this inhibition due to its enhanced ability to bind nucleosomes.

Surprisingly, our HMT assay results showed that PRC2-EZH1^Q571R^ could methylate H3K27 on H3K36me2-modified mono-nucleosomes to a degree similar to PRC2-EZH1^WT^ activity on unmodified mono-nucleosomes (Figure 5A). This suggests that PRC2-EZH1^Q571R^ might establish H3K27me3 domains in H3K36me2-enriched euchromatic regions. A similar trend was also observed with H3K36me3-modified mono-nucleosomes (Figure S5A).

**Figure 5.**
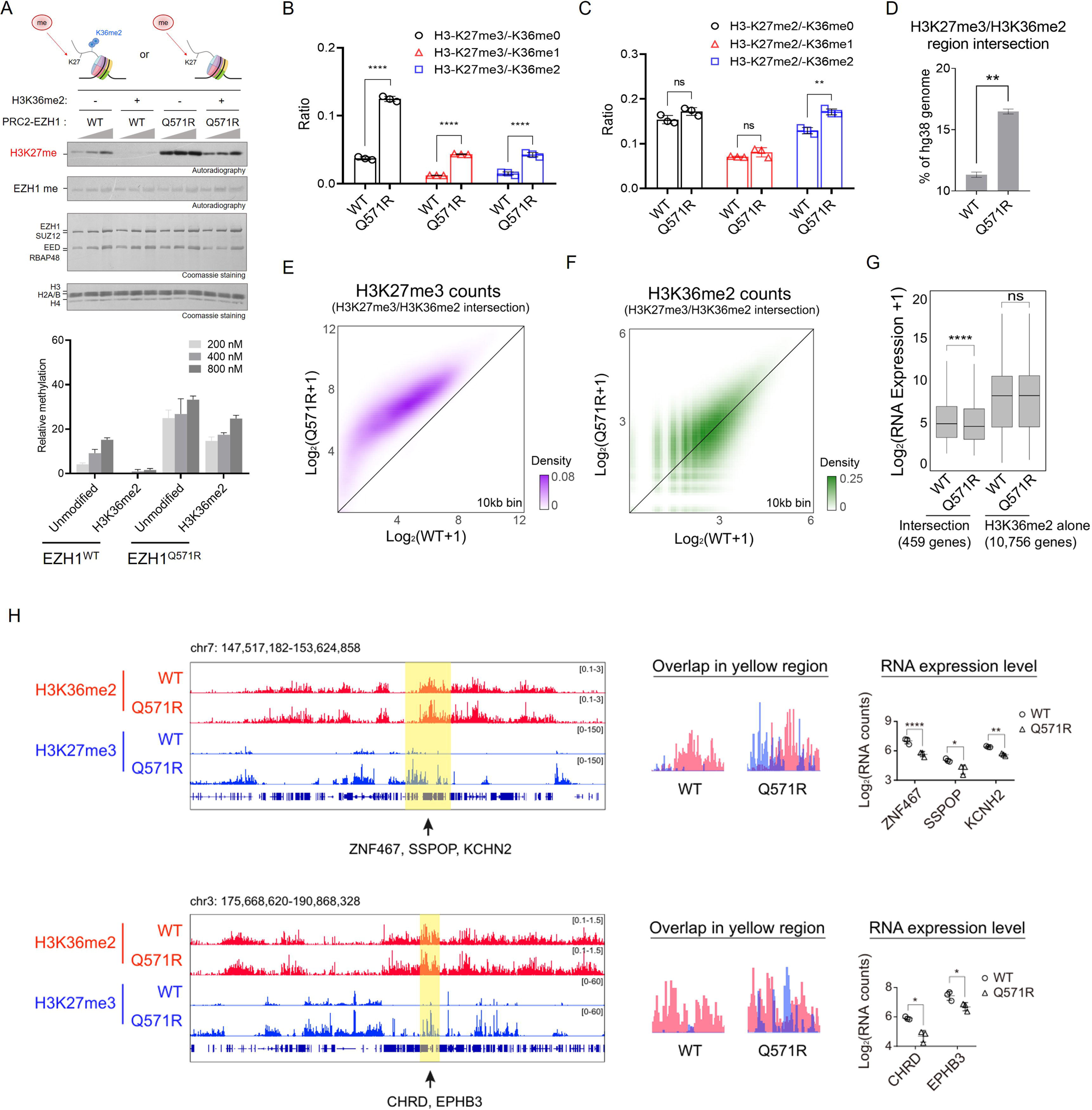
PRC2-EZH1^Q571R^ alters actively transcribed regions marked by H3K36me2/3. **(A)** Histone methyltransferase (HMT) assays. Data are presented as mean ± SEM, n=3 independent experiments. Top: HMT assay was conducted in the presence of 1 ug either unmodified or H3K36me2-modified mono-nucleosomes and varying concentrations of PRC2 (200, 400, or 800 nM). Incorporation of [^3^H]-SAM amounts into histone H3 or EZH1 was measured by autoradiography (top two images) and relative levels of PRC2 components (middle image) or mono-nucleosomes (bottom image) by Coomassie blue staining of SDS-PAGE gels. Bottom: Quantification of the relative level of [^3^H]-SAM incorporated into unmodified or H3K36me2-modified mono-nucleosomes. **(B-C)** Bar graphs show the relative ratios of H3K27me3 (B) or H3K27me2 (C) combined with different H3K36 methylation states (me0, me1, me2) in Nthy cells expressing EZH1^WT^ or EZH1^Q571R^. Data represent mean ± SEM (n=3). Welch’s t-test was used for statistical analysis (\*\*\**p*<0.001, \*\*\*\**p*<0.0001, \*\**p*<0.01, ns: not significant). **(D)** Bar graph showing the proportion of the hg38 genome represented by the intersection of H3K27me3 and H3K36me2 peak regions in Nthy cells expressing EZH1^WT^ or EZH1^Q571R^. Statistical differences were assessed using two-tailed Welch’s t-test (\*\**p*<0.01). **(E-F)** Scatter density plot showing the normalized H3K27me3 (E) or H3K36me2 (F) counts for each 10 kb bin at genomic regions identified by both H3K27me3 and H3K36me2 peaks in Nthy cells expressing EZH1^WT^ or EZH1^Q571R^. **(G)** Box plot showing the normalized RNA expression of genes located near H3K36me2 peak regions with (459 genes) or without (10,756 genes) significant upregulation of H3K27me3 in Nthy cells expressing EZH1^WT^ or EZH1^Q571R^. The significance threshold was set at *p*-value < 0.05 and log_2_ |fold change| > 1. Statistical differences were assessed using one-tailed paired *t*-test (\*\*\*\**p*<0.0001 and ns: not significant). **(H)** Left: Representative Integrative Genome Viewer (IGV) tracks showing the signals for H3K36me2 (red) and H3K27me3 (blue) in Nthy cell lines expressing EZH1^WT^ or EZH1^Q571R^. Yellow regions indicate the spread of H3K27me3 into actively transcribed regions marked by H3K36me2 in EZH1^Q571R^-expressing cells. Right: Zoomed-in views of yellow region and dot plots showing the log_2_(normalized RNA counts) for five genes located within yellow regions. Data are presented as mean ± SEM, n=3 independent experiments. Statistical differences were assessed using DESeq2 (\**p*<0.05, \*\**p*<0.01, and \*\*\*\**p*<0.0001).

To directly compare H3K27me2/3 and H3K36me2 modifications on the molecular level, we extracted histone proteins from EZH1^WT^ and EZH1^Q571R^-expressing Nthy cells and used mass spectrometry to quantify H3K36me2 in the presence of H3K27me2/3 within the same histone molecule. This revealed H3K27me2/3 and H3K36me2 to co-exist more frequently in EZH1^Q571R^-expressing cells (Figures 5B and 5C), indicating the ability of the mutant to catalyze H3K27 methylation in active chromatin regions.

Next, we conducted H3K36me2 and H3K27me3 CUT&Tag analysis to examine the coexistence and spread of H3K27me3 into active chromatin regions marked by H3K36me2. The number of genomic regions in which both H3K27me3 and H3K36me2 peaks were identified increased approximately 1.3-fold in EZH1^Q571R^-expressing cells (Figures 5D and S5B). This expansion was primarily driven by elevation of H3K27me3 within regions actively marked by H3K36me2 (Figure 5E), without significant alteration of H3K36me2 levels (Figure 5F). Notably, genes located near H3K36me2 peaks that acquired increased H3K27me3 (459 genes) exhibited significant transcript downregulation, whereas no significant change in RNA expression was observed across the total set of genes near H3K36me2 peak regions (10,756 genes) (Figure 5G). These results indicate that in the presence of EZH1^Q571R^, H3K27me3 intrudes into H3K36me2-enriched regions, repressing transcription of nearby genes (Figure 5H).

### PRC2-EZH1^Q571R^ exhibits a more pronounced elevation in HMT activity compared to PRC2-EZH2^Q570R^

Next, we sought to understand why Q to R mutation in the cancer context is frequently observed only in EZH1 and not in its counterpart, EZH2. Recent cryo-EM structures have revealed that in EZH1, the surrounding residues T565 and R562 are in close proximity to nucleosomal DNA, whereas the corresponding residues of EZH2 are more distantly situated (Figure 6A). This suggests that surrounding residues of EZH1 may contribute to the enhanced nucleosome binding affinity of the PRC2-EZH1^Q571R^ complex. Accordingly, we postulated that PRC2-EZH2 might demonstrate a lesser increase in HMT activity upon the EZH2^Q570R^ mutation. Supporting this idea, HMT activity assays did display an upsurge in activity for PRC2-EZH2^Q570R^, but significantly less than was observed for PRC2-EZH1^Q571R^ (Figure 6B). Specifically, the fold change in HMT activity relative to the corresponding wild-type form was 36.3 times greater for PRC2-EZH1^Q571R^ than for PRC2-EZH2^Q570R^ (Figure 6C). Moreover, the HMT activity of PRC2-EZH1^Q571R^ was significantly higher than that of PRC2-EZH2^WT^, indicating that the EZH1 mutant surpasses the strong catalytic activity of PRC2-EZH2.

**Figure 6.**
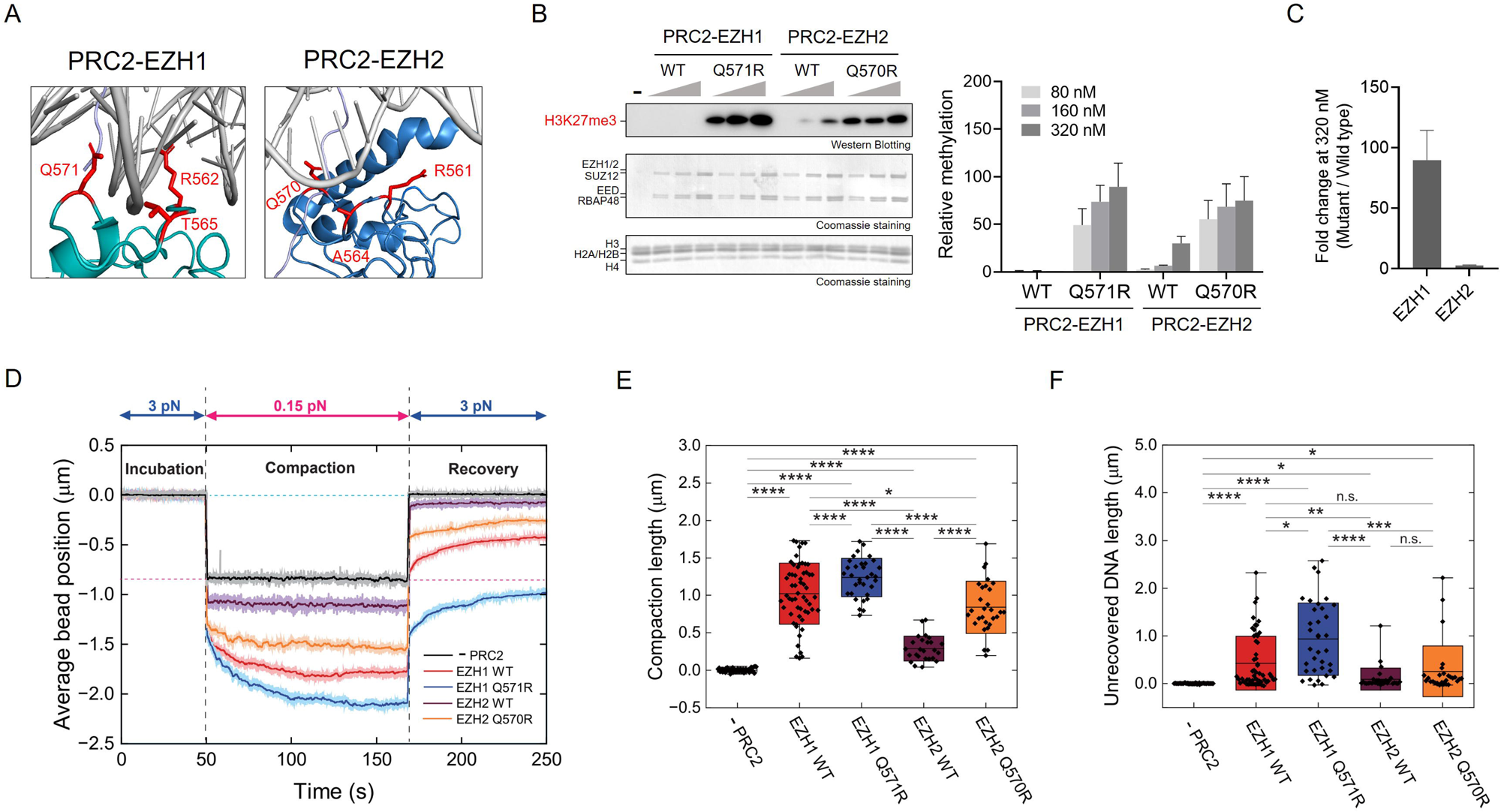
HMT and DNA compaction activities are more greatly enhanced for PRC2-EZH1^Q571R^ than for PRC2-EZH2^Q570R^. **(A)** Left: Closeup cryo-EM structure (modified from 7KTP and 7KTQ) showing interaction of the EZH1 CXC domain (sky blue) with nucleosomal DNA (gray). R562, T565, and Q571 residues are indicated in red. Right: Closeup cryo-EM structure (modified from 6WKR) showing interaction of the EZH2 CXC domain (blue) with nucleosomal DNA (gray). R561, A564, and Q570 residues are indicated in red. **(B)** Histone methyltransferase (HMT) assays. Data are presented as mean ± SEM, n=3 independent experiments. Left: HMT assay was conducted in the presence of 300 ng di-nucleosome and varying concentrations of PRC2 (80, 160, or 320 nM). H3K27me3 was measured by western blot (top image) and the relative concentration of PRC2 components (middle image) or di-nucleosomes (bottom image) by Coomassie blue staining of SDS-PAGE gels. Right: Quantification of the relative level of H3K27me3-modified di-nucleosomes. **(C)** Bar graph showing the fold change in HMT activity (mutant / wild type) observed when using 320 nM PRC2-EZH1 or PRC2-EZH2. Data are presented as mean ± SEM, n=3 independent experiments. **(D)** Average trace of DNA compaction and recovery for PRC2 with wild-type and mutant forms of EZH1 and EZH2. Thick lines with full color opacity represent moving averages calculated over 100 points, while lower-opacity halos represent the raw data. **(E-F)** Plots of DNA compaction length (E) and unrecovered DNA length (C) for various complex conditions. N = 57 for no PRC2 and 57, 35, 25, and 29 for, PRC2 possessing EZH1 WT, EZH1 Q571R, EZH2 wildtype, and EZH2 Q570R, respectively. Box plots represent the range from mean – s.d. to mean + s.d.; center lines show means and whiskers the minimum and maximum values. *P*-values were assessed using a two-tailed Student’s *t*-test (\**p*<0.05, \*\**p*<0.01, \*\*\**p*<0.001, \*\*\*\**p*<0.0001, and ns: not significant).

We further utilized magnetic tweezer analysis to examine the ability of PRC2-EZH2^Q570R^ to maintain DNA compaction compared to PRC2-EZH1^Q571R^ and the corresponding wild-type complexes. Determination of average bead positions revealed PRC2-EZH2 to more weakly interact with DNA than PRC2-EZH1 (Figure 6D). This observation aligns with previous findings indicating that PRC2-EZH1 promotes PRC2/DNA clustering, whereas PRC2-EZH2 binds to DNA without forming clusters.^8^ PRC2-EZH2^WT^ exhibited initial compaction but showed minimal further compaction at 0.15 pN (Figure 6E). In contrast, the PRC2-EZH2^Q570R^ mutant displayed greater compaction, resembling the behavior of PRC2-EZH1^WT^, and the PRC2-EZH1^Q571R^ mutant exhibited still further enhanced compaction. However, the increase in unrecovered DNA length relative to wild-type was significantly greater for mutant PRC2-EZH1 (Figure 6F), suggesting that only PRC2-EZH1^Q571R^, not PRC2-EZH2^Q570R^, induces irreversible DNA compaction.

## Discussion

Recently, the development and clinical trial of EZH1/2 dual inhibitors has led EZH1 to garner significant interest in cancer research,^39,40^ and consequently to an essential inquiry into the unique roles of EZH1 and EZH2 in the cancer context. In our study, we aimed to shed light on the specific mechanism of chromatin compaction associated with EZH1 by closely examining the EZH1^Q571R^ mutant, which occurs naturally in thyroid cancer.

Compared to PRC2-EZH2, PRC2-EZH1 exhibits stronger nucleosome and DNA binding,^5,7,8^ primarily due to enrichment of positively charged residues in its SANT1 and SANT2 domains. In this study, we further identified a single residue within the CXC domain that contribute to enhanced nucleosomal DNA interactions. Structural analysis of PRC2-EZH1 with mono-nucleosomes (Figures 3A and 3B) revealed a critical interaction between the Q571 residue of EZH1 and nucleosomal DNA. Notably, in the dimeric form of PRC2-EZH1, two Q571 residues are positioned at the nucleosomal exit site.^8^ In the Q571R mutation, the replacement of glutamine (Q) with arginine (R) strengthens electrostatic interactions, resulting in a highly stable and compact chromatin structure that could effectively lock the nucleosome and prevent DNA unwrapping. Interestingly, a Q571K mutation, which introduces another positively charged residue, has also been identified in thyroid cancer (Figure 1A), reinforcing the idea that positively charged residues play crucial roles in nucleosomal DNA binding. Furthermore, we confirmed that the PRC2 complex containing the EZH1 Q571R mutation specifically enhances methyltransferase activity toward histones within chromatin, but not toward non-histone substrates such as EZH1 itself.

The chromatin compaction mechanism of PRC2-EZH1 may be similar to but also distinct from other well-characterized pathways, such as HP1- and PRC1-mediated compaction. HP1 bridges neighboring nucleosomes via H3K9me3 binding, while PRC1 utilizes CBX subunits and the PHC-SAM domain to form phase condensates, promoting chromatin compaction.^41–45^ Interestingly, PRC2-EZH1 forms condensates more efficiently than PRC2-EZH2,^5^ hinting at it having a role in polycomb-mediated chromatin condensation along with PRC1. Unlike HP1 and PRC1, EZH1 serves as both a histone methyltransferase (H3K27me3 writer) and a chromatin compactor, suggesting a unique dual function. Further investigation is needed to determine whether EZH1 acts as a prerequisite for PRC1-PHC in chromatin compaction, particularly in differentiated cells.

A key question remains: why is the EZH1^Q571R^ mutation specifically found in thyroid cancer? One possible explanation is that normal follicular thyroid epithelial cells are highly differentiated, leading to relatively high EZH1 expression compared to undifferentiated cells. The EZH1^Q571R^ mutation may trigger a transition to a proliferative state, contributing to tumorigenesis. Notably, the PRC2-EZH1^Q571R^ mutant exhibits significantly higher activity than wild-type PRC2-EZH2 (Figure 6B), and may thereby mimic the effects of the EZH2 overexpression or gain-of-function mutations commonly observed in many cancers. Additionally, EZH1^Q571R^ is strongly linked to the cAMP pathway and activating mutations in TSHR,^22^ both of which play central roles in thyroid tumorigenesis. This association provides a clear mechanistic link between EZH1 mutation and thyroid cancer. Notably, although EZH1^Q571R^ is predominantly found in thyroid cancer, a different substitution at the same residue (Q571L) has been identified in non-small cell lung cancer, a differentiated epithelial cancer.^46^ Whether EZH1 mutations are present in other differentiated-cell-derived cancers, such as neuronal tumors, remains an open question. It merits mention that EZH1 gain-of-function mutations are also frequently observed in neurodevelopmental disorders,^47^ albeit outside of a cancerous context.

Another key finding is that PRC2-EZH1^Q571R^ efficiently catalyzes H3K27me2 and H3K27me3 on pre-existing H3K36me2 within the cell. While PRC2-EZH1^Q571R^ is also capable of catalyzing H3K27me3 on pre-existing H3K36me3 *in vitro*, co-existence of H3K27me3 and H3K36me3 was not detected in cells through mass spectrometry analysis. This absence may be due to the extremely low co-occurrence of H3K27me3 and H3K36me3.

The antagonistic relationship between H3K27me3 and H3K36me2/3 has been extensively documented, and its dysregulation is linked to tumorigenesis, including of diffuse intrinsic pontine glioma.^48^ Structural studies have provided insight into this antagonism, with one particularly relevant work showing that H3K36me3 occupies a position that interfaces with EZH2^Q570^ and nucleosomal DNA.^38^ Interestingly, the unmethylated form of H3K36 does not occupy the same site; instead, this positioning occurs specifically upon trimethylation (Figure S5C). Thus, EZH1^Q571R^ may outcompete H3K36me2/3 due to its stronger affinity for nucleosomal DNA, which aligns with our HMT assay and epigenome data (Figures 5A, 5D-H and S5A). Previous studies have proposed that H3K27me3-heterochromatin can invade euchromatic regions through the activity of H3K36 demethylases, which remove H3K36 methylation, thereby enabling subsequent H3K27 methylation. In contrast, our study demonstrates that PRC2-EZH1^Q571R^ can independently invade euchromatin both *in vitro* and within cells.

Lastly, how should we approach the treatment of patients with thyroid cancer expressing EZH1^Q571R^? While clinically approved PRC2 active site inhibitors are available, the increased HMT activity of EZH1^Q571R^ is due to the enhanced interaction of EZH1^Q571R^ with nucleosomal DNA that augments DNA compaction. In addition, a recent study has shown that K562 cell lines expressing EZH2^Q570R^ exhibit resistance to the PRC2-EZH2 active site inhibitor GSK 343.^49^ This underscores the need to develop a novel inhibitor targeting the EZH1^Q571R^-nucleosomal DNA interface. One potential strategy involves the development of synthetic RNA molecules designed to induce aberrant PRC2 dimerization, which renders it inactive. A recent study has demonstrated that G-quadruplex RNA, targeting EZH2 353-362, inhibits PRC2 activity by promoting PRC2 dimerization, and that this dimerization process is mediated by interactions between the individual EZH2 CXC domains, including residues R561, K563, T568, and Q570.^50^ Hence, synthetic RNA molecules have the potential to inhibit the increased PRC2 activity by disrupting the interaction between EZH1^Q571R^ and nucleosomal DNA. Given that hyperactive EZH1 mutations are also found in neurodevelopmental diseases,^47^ developing an EZH1-specific inhibitor could have therapeutic applications for both cancer and neurodevelopmental diseases.

## Resource Availability

### Lead contact

Further information and requests for resources and reagents should be directed to and will be fulfilled by the lead contact, Chul-Hwan Lee.

### Materials availability

Materials are available from Chul-Hwan Lee upon request.

### Data and code availability

The NGS data reported in this study have been deposited in the GEO repository on the NCBI website GSE294320 for ATAC-seq, GSE294321 for EZH1 ChIP-seq, GSE294323 for H3K27me3 and H3K36me2 CUT&Tag, and GSE294325 for RNA-seq.

This paper does not report original code.

Any additional information required to reanalyze the data reported in this paper is available from the lead contact upon request.

## Acknowledgements

We thank Dr. Jia-Ray Yu for critical reading of the manuscript as well as Lee laboratory members for critical comments and discussions. This study was supported by the Institute of Information & Communications Technology Planning & Evaluation (IITP) grant, funded by the Korean government (MSIT) (grant numbers RS-2023-00223069). Also, it was supported by the National Research Foundation of Korea (grant numbers NRF2021R1C1C1013220 and NRF2022R1A5A102641311) and the BK21 Four Biomedical Science Program. The SNUH Kun-hee Lee Child Cancer and Rare Disease Project Foundation, Republic of Korea (grant number 22B-001-0100), the Research Resettlement Fund for the new faculty of Seoul National University, the Creative-Pioneering Researchers Program through Seoul National University, grants from Seoul National University College of Medicine, the AI-Bio Research Grant through Seoul National University also supported this study. Furthermore, it was supported by Doosan Yonkang Foundation (Grant No. 30-2024-0440) and by NAVER. The figures were created with BioRender.com. The results shown in Figures 2 and S2 are in part based upon data generated by the TCGA Research Network (https://www.cancer.gov/tcga) and the GTEx Portal (https://www.gtexportal.org/home/) on 01/10/2025. J.-K.R. acknowledges the Institute of Applied Physics of Seoul National University, Creative-Pioneering Researchers Program and the Research Grant (3348-20220014, 370C-20230124) from Seoul National University, the Brain Korea 21 Four Project grant funded by the Korean Ministry of Education, and the National Research Foundation of Korea (Project Number RS-2023-00212694, RS-2023-00265412, RS-2023-00218318, RS-2022-NR066591, RS-2024-00410733 and RS-2023-00301976).

## Author contributions

H.K., D-G.K., S.H., K.E.L., J-K.R. and C-H.L. conceptualized and designed the study. H.K., D-G.K., S.H., J-Y.J., Y.O., and H.K. conducted the experiments. F.N.L.V. and J.M.G performed quantitative mass spectrometry analyses in B.A.G.’s lab. H.K. in C-H.L. lab and G.P in I.J. lab performed bioinformatics analyses. S-H.I, C-K.J. and S-Y.K. analyzed EZH1 mutations found in thyroid cancer. W.K., J-K.W., S.K and K.E.L prepared thyroid tissue samples. H.K., D-G.K., S.H., K.E.L., J-K.R. and C-H.L. wrote the manuscript.

## Declaration of generative AI and AI-assisted technologies

No generative AI or AI-assisted technologies were used in the preparation of this work.

## Supplemental information

Document S1. Figures S1-S5 and Table S1

## Materials and Methods

### Protein purification using a baculovirus expression system

To purify human PRC2 core complexes, FLAG-EED, EZH1, SUZ12, and RBAP48 were cloned independently into a pFASTBac1 vector (Invitrogen,10-360-014). EZH1 mutant constructs were generated by site-directed mutagenesis and mutations were confirmed by Sanger DNA sequencing. All four core components of PRC2 (FLAG-EED, EZH1 or EZH2, SUZ12, and RBAP48) were co-expressed in Sf9 cells by baculovirus infection. After 60 hours of infection, Sf9 cells were resuspended in BC350 buffer (20 mM HEPES-NaOH, pH 7.8, 1 mM EDTA, 350 mM NaCl, 10% glycerol, and 0.1% NP-40) with protease inhibitors (1 mM phenylmethylsulfonyl fluoride, 1 mM benzamidine, 5 µg/ml leupeptin and 5 µg/ml pepstatin A) and phosphatase inhibitors (10 mM NaF and 1 mM Na_3_VO_4_). Cells were lysed by sonication (Sonics Vibra Cell VCX-130), and WT or mutant recombinant PRC2 was purified by FLAG-M2 agarose beads (Sigma, A2220) and Q Sepharose beads (Cytiva, 17051010).

### Nucleosome reconstitution

Recombinant histones were generated as previously described.^51^ Briefly, each core histone was expressed in Rosetta (DE3) cells (Enzynomics, CP-1011), extracted from inclusion bodies, and purified by HiTrap SP sepharose FF (Cytiva, 17505401). To refold the recombinant octamers, equal amounts of histones were combined and dialyzed into a refolding buffer (10 mM Tris-HCl, pH 7.5, 2 M NaCl, 1 mM EDTA, and 5 mM β-mercaptoethanol). Octamers were further purified using size exclusion chromatography on a 24-mL Superdex 200 column (Cytiva, 28990944). Recombinant nucleosomes were reconstituted by sequential salt dilution of octamers and plasmid having 601 nucleosome positioning sequences. Recombinant mono-nucleosomes modified with H3K36me2 (Epicypher, 16-0319) or H3K36me3 (Epicypher, 16-1320) and paired unmodified mono-nucleosomes (Epicypher, 16-0009) were purchased from Epicypher.

### Histone Methyl Transferase (HMT) assay

Standard HMT assays were performed in a 15 µL reaction mixture consisting of HMT buffer (50 mM Tris-HCl, pH 8.5, 5 mM MgCl_2_, and 4 mM DTT), 50 µM of S-Adenosylmethionine (SAM) (Sigma, A7007), 300 ng of nucleosomes, and recombinant human PRC2 complexes. The mixtures were incubated at 30℃ for 60 minutes and then stopped by adding 3.75 µL of lammeli buffer (300 mM Tris-HCl, pH 6.8, 50% glycerol, 10% SDS, 25% β-mercaptoethanol, and 0.01% bromophenol blue). The extent of methylation of each HMT reaction was measured using the following steps: after HMT reactions, samples were incubated at 95℃ for 5 minutes, separated on SDS-PAGE gels, transferred to 0.45 um PVDF membranes (Millipore, IPVH00010) and then probed using an H3K27me3 antibody (CST, 9733). When [³H]-SAM (Revvity, NET155V250UC) was used in the HMT assay, the PVDF membrane was exposed to X-ray film.

### Electromobility shift assay

Recombinant PRC2 complexes and mononucleosomes were dialyzed into EMSA binding buffer (EBB) (10 mM HEPES, pH 7.9, 250 mM KCl, 5% v/v glycerol, 5 mM DTT, and 0.5 mg/ml BSA). PRC2 proteins were serially diluted in EBB, and an equal volume of 80 ng nucleosome containing 10 µM SAM was then added to each tube. The mixtures were incubated at 30℃ for 45 minutes and then loaded onto 3.5% native polyacrylamide gels. The gels were run at 120 V for 70 minutes using 0.3× Tris Borate EDTA running buffer (30 mM Tris base, 30 mM boric acid, and 0.6 mM EDTA). Subsequently, the gels were stained using SYBR Gold (ThermoFisher, S11494) and scanned using the ChemiDoc MP System (Bio-rad).

### Preparation of non-coilable biotin-labeled **λ**-DNA

Unmethylated circular λ-DNA (Promega, N3011S) was modified with Taq ligase (NEB, M0208L), along with complementary oligomers JT41 (5’-phosphate-GGGCGGCGACCT-biotin-3’) and JT42 (5’-phosphate-AGGTCGCCGCCC-biotin-3’) (Integrated DNA Technologies). The resulting biotin-labeled DNA was purified using AKTA pure system (Cytiva) and size exclusion chromatography was performed using Sephacryl S-500 HR resin (Cytiva, 17061310).

### Single-molecule fluorescence assay

We followed a well-established single-molecule fluorescence assay protocol (Figures 4 and S4).^33^ To prepare the assay, we drilled 12 holes at the side of glass slides to create input and output channels. We cleaned the slides with 10% detergent, acetone, Mili-Q water, and Piranha solution and then polyethylene glycol (PEG)ylated them with a 1:80 ratio of biotin-PEG and PEG by incubating them for 12-24 hours in a sodium bicarbonate solution (0.1 M NaHCO_3_, pH 8.5). After washing the slides with Mili-Q water and drying them with nitrogen gas, we carried out additional PEGylation with MS(PEG) solution. The same process is executed as before. The only difference is incubating time (1-24 hours). Last, we partitioned the channels with double-sticky tapes and sealed the edges with epoxy glue.

To immobilize 48.5-kbp length of double-biotinylated λ-DNA, we flowed streptavidin (100 µg/ml) in T50 buffer (40 mM Tris, pH 7.5 and 50 mM NaCl) through the channels and incubated for 2 minutes to induce streptavidin biotin-PEG binding. We then washed the channels with T50 buffer, and we applied 48.5-kbp length of double-biotinylated λ-DNA in imaging buffer (40 mM Tris, 100 mM NaCl, 0.5 mM TCEP, 0.25 mg/ml BSA, and 50 nM Sytox Orange (SxO) (Thermofisher, S11368) at a flow speed of 8 µl/m. After DNA tethering, we removed unbound DNA by flowing the imaging buffer. To monitor PRC2-induced DNA compaction, we flowed the PRC2 solution (50 nM PRC2 with imaging buffer) through the channels at a speed of 20 µl/m for 1 minute. To image the samples, we used a 1mW 561-nm HILO mode laser for excitation of SxO. Using an Olympus UPlanXApo 100x /1.45 lens and an Andor iXon Ultra 897 EM CCD, we imaged real-time DNA compaction.

### Single-molecule analysis

To obtain kymographs, the intensity values of *n* pixels (*n* is proportional to the region of interest (ROI)) from the ROI perpendicular to the extended DNA were summed in each frame. Although the appearance of clusters was sometimes not obvious due to the resolution limit, we determined the compaction time using a fluctuation analysis (Figures S4A and S4B). Specifically, as the cluster size (or number of clusters) increased, the fluctuation decreased. The term “fluctuation” referred to the variance of the intensity region between adjacent frames (Figures S4A and S4B).

To quantify the fluctuation, the mean of the intensity values below the 50th percentile within the DNA ROI was taken as the background noise and subtracted from the DNA signal. To focus on the DNA variance, the background intensity was set to zero (All of the ‘black (background)’ in the DNA figures indicates zero intensity). To ensure that each DNA had the same length in every frame, the sum of the DNA signal was equalized across all frames after background removal. The overlapped region between the present and previous frames (Figure S4A) was compared, and only the variance parts (Figure S4B) were used to calculate the sum of the variance intensity, which was defined as the fluctuation previously.

To measure the compaction time, the fluctuation data was smoothed using Savitzky-Golay with a moving window of 150 or 300 and a polynomial order of 2, and then normalized. Specifically, the starting point was set at the point where the DNA signal showed a large fluctuation that soon decreased, indicating the onset of compaction. Finally, exponential decay equation 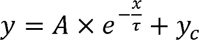 was fitted to the fluctuation data to determine the compaction time.

### DNA construction for magnetic tweezers experiment

DNA was constructed by joint two DNA fragments for the magnetic tweezers experiment (Figures 4 and 6). The first fragment, referred to as the main template, was amplified by PCR from the pBIG1a plasmid (sequences in the supplementary information). The PCR was performed using the following primers 5’-biotin-GACTGCAGGCTCTAGACTACT-3’ and 5’-CCGGAATTCCGGACTGGCCTCAGGCATTTGAG-3’. The second fragment, termed the handle, was generated by PCR using the primers 5’-CCGGAATTCCGGCAGGCTCTAGACTACTGCTCA-3’ and 5’-GTGTGCTTCAGCGTATTGCC-3’. The PCR reaction for this fragment included a 1:5 ratio of dTTP to dUTP-11-DIG (Jena Bioscience, NU-803-DIGX-L). Following PCR, both fragments were purified using a PCR purification kit (Qiagen, 28104). The purified fragments were mixed in equimolar concentrations and subsequently digested with the EcoR1-HF restriction enzyme (NEB, R3101S). The fragments and mixed 1:1 about mol concentration. And they digested with EcoR1-HF restriction enzyme and washed with PCR purification kit. The resulting mixture was used for ligation with T4 ligase (Enzynomics, M001S). After the ligation step, the DNA sample was loaded onto an agarose gel, purified through electrophoresis, and extracted using a gel extraction kit (Qiagen, 28704). By doing so, the two DNA fragments were joined to construct a 9.2 kbp length of DNA molecule.

### Magnetic Tweezers Setup

We used a magnetic-tweezers setup built by following previous literature.^35^ In brief, a linear stage motor (Physik Instrument, M-126.PD2) was utilized to control the movement of the magnets, thereby adjusting the magnetic field applied to the magnetic beads tethered to DNA molecules. The beads were illuminated using a green LED light (Mouser Electronics, 941-XPEBGRL100M01). The diffraction pattern from the beads was collected through a CFI Plan 50XH objective (50x magnification, NA=0.9, Nikon) and recorded at 90 Hz using a CMOS camera (Teledyne Dalsa, Falcon2 12M FA-80-12M1H). Data acquisition was conducted with a LabView program and data analysis were performed using custom Python scripts.

### Magnetic tweezers flow-cell Preparation

The flow cell chamber was constructed as described previously.^35^ The chamber was assembled by overlapping two cover slips with a spacer tape placed between them. The bottom coverslip (VWR, 24 x 60 mm^2^, thickness No. 1.5, 48393-251) served as the imaging surface to which the DNA molecules were attached. The top coverslip (VWR, 24 x 50 mm^2^, thickness No 1.5, 48393-241) was placed over the bottom coverslip, creating a flow channel between the two slides through which buffer and protein samples could be passed. Before assembly, the bottom coverslip was sonicated in distilled water (DW) for 5 minutes, followed by three times of washing with DW. It was then sonicated in 1 M KOH for 20 minutes, followed by three additional washes with DW. After a final 5-minute sonication in DW and three additional DW washes, the coverslip was dried using nitrogen gas. Once the bottom coverslip was completely dried, 10 μL of 100x diluted polystyrene beads (Polybead, 17133-15) were coated onto the coverslip using the lateral side of a 10 μL pipet tip. The coverslip was then heated on a hot plate at 150°C for 50 seconds to firmly attach the polystyrene beads, which served as reference beads. Following this, the bottom slide was coated with a 0.1% nitrocellulose membrane (Invitrogen, LC2001) solution in pentyl acetate (Sigma Aldrich, 109584), prepared by diluting a 1% nitrocellulose solution in pentyl acetate.

After the pentyl acetate evaporated, the bottom coverslip was placed on a flat surface for flow cell assembly. Double-sided sticky tape (Scotch 3M, 237ROK) was attached to the sides of the bottom coverslip, creating a narrow flow path. The top coverslip was sonicated in DW for 5 minutes, followed by three DW washes. It was then sonicated in a solution of 70% acetone and 30% isopropanol, followed by three additional DW washes. The coverslip was then dried using nitrogen gas. The top coverslip was carefully placed on the bottom coverslip, ensuring alignment and creating a space for the flow channel. Pipet tips were used to create both the injection and output ports at the ends of the flow cell. Small openside were made at each end, where pipet tips were securely attached using epoxy glue (Devcon, 20845), ensuring a leak-proof seal between the pipet tips and the coverslip surface. On the injection side, a pipet tip was attached and cut at the wider end to serve as a reservoir. This reservoir allowed for the introduction of DNA, buffer, or protein samples into the flow cell. On the opposite side, another pipet tip was attached in the same manner but was additionally connected to a piece of flexible tubing. This tubing was linked to a syringe system, enabling controlled suction. This setup allowed precise control of flow rates within the chamber, ensuring that samples moved steadily through the flow cell without causing turbulence. The assembled flow cell was incubated in a solution of 100 μg/mL anti-digoxigenin (Roche, 11333089001) in PBS buffer, pH 7.5 (Enzynomics, EBP006-1000) for 1 hour. After incubation, the flow cell was washed with 1 mL of PBS containing 1% BSA (Sigma Aldrich, A7906), followed by an overnight incubation in PBS/BSA buffer.

### Magnetic-tweezers experiment

A 9.2 kbp length of DNA fragment was diluted to a final concentration of 16 pM in PBS/BSA 1% buffer. A 100 μL aliquot of the diluted DNA solution was introduced into the flow cell and incubated for 1 hour to allow the DNA to attach to the bottom coverslip. The flow cell was then washed with PBS/BSA buffer to remove any unbound DNA. The flow cell was mounted on the magnetic tweezers, and the reference beads were located using the microscope. The magnet height was adjusted to 18 mm. A 40 μL solution of 10x diluted MyOne magnetic beads (Invitrogen, 65001) was introduced into the flow cell at a flow rate of 10 μL/min. After bead loading, the flow cell was washed again with PBS/BSA buffer, followed by a wash with PRC2 salt buffer (1 M NaCl, 0.1 M Tris-HCl, pH 7.5, and 0.5 mM TCEP (Millipore, 580561)) and equilibrate with the PRC2 reaction buffer (50 mM NaCl, 0.1 M Tris-HCl, pH 7.5, and 0.5 mM TCEP). We applied 10 nM PRC2 proteins with various mutants in EZH1 or EZH2 in the PRC2 reaction buffer. A total of 200 μL of this protein solution was injected into the flow cell at a flow rate of 100 μL/min applying with strong magnet force 3 pN. After injection, a waiting time of 50 seconds was allowed to stabilize the flow rate and minimize turbulence in the chamber. After the flow rate stabilized, magnetic force measurements were conducted. The magnetic force was applied in a sequence as follows: 50 seconds at 3 pN, 120 seconds at 0.15 pN, and 100 seconds at 3 pN. Changing force, magnetic is vertically moving with 2 mm/s and it takes around 1.46 seconds for force transition. We observed rapid DNA compaction when the force was decreased to 0.15 pN. Data were recorded throughout the experiment and analyzed using custom Python scripts.

### Cell culture and generation of isogenic cell line

Nthy cells were cultured in standard RPMI-1640 medium (Biowest, L0498) supplemented with 10% FBS (Biowest, 51480), 2 mM GlutaMax (ThermoFisher, 35050-061), and 1% antibiotics/antimycotics (Biowest, L0010). Isogenic cell lines that constitutively expressing EZH1 WT or Q571R were generated by a lentiviral delivery system. After lentiviral infection, Nthy cells were sorted using BD AriaIII based on mCherry expression from the pLV-tetO-mCherry vector (Addgene, 70273).

### Lentiviral production and delivery

The pLV-tetO-mCherry vector encoding EZH1 WT or Q571R was prepared for lentiviral production and delivery. For the production of viral particles, each 10 µg of lentiviral vector, packaging vectors (2.5 µg of pcREV, 3 µg of BH-10, and 5 µg of pVSV-G) was transfected using PEI (Merk, 919012) into Lenti-X 293T cells. The virus-containing medium was collected 48 hr after transfection and added to the target cells in the presence of 8 ng/mL polybrene (Merk, TR-1003G). Infected Nthy cells were sorted using a BD AriaIII after 2 days.

### Proliferation assay

2×10^5^ Nthy-ori-3-1 cells that constitutively expressing empty vector, EZH1^WT^ or EZH1^Q571R^ were seeded onto 10-cm plates. After 48 hours, cells were detached using a 0.25% Trypsin-EDTA solution (Gibco, 25200072) and cell numbers were measured by hemocytometer.

### Preparation of whole cell extract and western blotting

A total of 8×10^6^ cells were harvested and lysed using RIPA buffer (10 mM Tris-HCl, pH 7.5, 150 mM NaCl, 1 mM EDTA, 1% Triton X-100, 0.1% sodium deoxycholate, and 0.1% SDS) supplemented with protease inhibitors (1 mM phenylmethylsulfonyl fluoride, 5 µg/mL pepstatin A, 5 µg/mL leupeptin, and 1 mM benzamidine) and phosphatase inhibitors (10 mM NaF and 1 mM Na_3_VO_4_). The lysates were sonicated using VCX-130 and centrifuged at 18,000 x g for 20 minutes at 4℃. The resulting supernatant was collected as the whole cell extract. For western blotting, samples were prepared by adding Laemmli buffer (2% SDS, 5% β-mercaptoethanol, 10% glycerol, 0.002% bromophenol blue, and 60 mM Tris-HCl, pH 6.8) to the whole cell extract. Samples were separated by 8-16% SDS-PAGE gels and were transferred to 0.45-µm PVDF membranes. The antibodies used are as follows; H3K27me3 (CST, 9733), H3K9me3 (Abcam, ab8898), H3K27me2 (CST, 9728), H3K27ac (Abcam, ab4729), H3 (Abcam, ab1791), EZH1 (Proteintech, 20852-1-AP), EZH2 (CST, 5246), SUZ12 (CST, 3737), β-tubulin (Abbkine, A01030), LMNA/C (CST, 4777), and GAPDH (GeneTex, GTX627408).

### Immunofluorescence

A total of 2×10⁴ cells were seeded onto a 2-well cell culture slide (SPL, 30402). After 24 hours, the cells were washed with 1x PBS and fixed with 4% paraformaldehyde (Biosesang, PC2031) for 20 minutes. Following fixation, the cells were permeabilized with a permeabilization buffer (1% BSA and 0.1% Triton X-100 in 1x PBS) for 20 minutes and then washed three times with 1x PBS. The primary antibody, diluted in antibody buffer (1% BSA in 1x PBS), was incubated overnight at 4°C. After three washes with 1x PBS, Alexa Fluor 647 secondary antibody (ThermoFisher, A32733), diluted in antibody buffer, was incubated for 1 hour at room temperature, followed by three additional washes with 1x PBS. Mounting medium was then applied dropwise to the cells, and a coverslip was placed over them. Imaging was performed using the LSM-980 (ZEISS).

### Subcellular protein fractionation

Subcellular protein fractionation was conducted using Subcellular Protein Fractionation Kit for Cultured Cells (Thermo Fisher, 78840) and analyzed using western blotting protocol.

### Histone extraction and mass spectrometry

Cells were collected using a scraper and harvested with 1x PBS (Enzynomics, EBP-001-500) by centrifuging at 800 xg for 5 minutes at room temperature. The pellet was then washed twice with 5 volumes of nuclear isolation wash buffer (NIB wash) (15 mM Tris-HCl, pH 7.5, 15 mM NaCl, 60 mM KCl, 5 mM MgCl₂, 1 mM CaCl₂, 250 mM Sucrose, 600 µM AEBSF, 1 mM DTT, and 10 mM sodium butyrate). Next, pellet was resuspended in 10 volumes of NIB wash with 0.3% NP-40 alternative (Sigma, 492018) and left on ice for 5 minutes. Nuclei were collected by centrifugation at 700 x g for 5 minutes at 4°C and washed three times with 10 volumes of NIB wash. The isolated chromatin was resuspended in 5 volumes of 0.2N H₂SO₄ and rotated for 2 hours at 4°C. Following centrifugation at 3400 x g for 5 minutes at 4°C, the supernatant was transferred to new tubes, and histones were precipitated overnight using 25% TCA (Duksan, 76-03-9). After centrifugation at 3400 xg for 5 minutes at 4°C, the histones were washed three times with cold acetone. The dried pellet was resuspended in distilled water and prepared for mass spectrometry analysis. Mass spectrometry and data analysis were conducted as previously described.^52,53^

### Genomic DNA extraction and Sanger sequencing

Thyroid tissues were resuspended with genomic DNA extraction buffer (50 mM Tris-HCl, pH 8.0, 2 mM NaCl, 10 mM EDTA, pH 8.0, and 0.1% SDS) and homogenized using a homogenizer. The lysate was then incubated at 55°C for 1 hour with proteinase K (Thermo Fisher, EO0491). After incubation, insoluble material was pelleted, and the supernatant was transferred to new tubes. Genomic DNA was subsequently purified using phenol/chloroform/isoamyl alcohol (Sigma, P3803). 50ng of genomic DNA was used for PCR to determine the nucleotide sequence of EZH1. The primer sequences used for PCR were as follows: forward primer (5’-ctgaagctgccaacagatgagctca-3’) and reverse primer (5’-atagacctttccgcgtcgatcagcc-3’).

### RT-qPCR

Thyroid normal and tumor tissues were resuspended in 1 ml of trizol (ThermoFisher, 15596026) and incubated for 5 minutes at room temperature. Then, 0.2 ml of Chloroform (Sigma, C2432) was added, and the solution was vortexed vigorously. The mixture was centrifuged at 13,000 rpm for 15 minutes at 4°C, and the upper phase was transferred to a fresh tube. RNA was precipitated using isopropanol (Duksan, 857) and washed with ice-cold 70% ethanol (Sigma, E7023). The precipitated RNA was dissolved in DEPC-treated water (Invitrogen, AM9915G), and RNA concentration was measured. 1 μg of RNA was used to synthesize cDNA using AccuPower RT Premix (Bioneer, K-2041). qPCR was performed using RbTaq SYBR Green qPCR PreMIX (Enzynomics, RT-530) and primers (see supplementary table).

### Ethics Statement

This study was approved by the Institutional Review Board of the Seoul National University Hospital (IRB No.: 0809-097-258). All procedures were conducted in accordance with the ethical standards of the institutional and/or national research committee and with the 1964 Helsinki Declaration and its later amendments or comparable ethical standards. Written informed consent was obtained from all individual participants included in the study.

### Patients and Specimen Collection

Fresh frozen thyroid tissue samples were collected from 74 patients (5 oncocytic thyroid cancers and 69 follicular thyroid cancers) who underwent thyroid surgery between December 2012 and August 2024. The final diagnosis of each case was determined based on histopathological evaluation of the thyroidectomy specimens. The presence or absence of capsular invasion was assessed by an experienced pathologist (J.K.W.) through microscopic examination of hematoxylin and eosin stained slides.

### ATAC-seq and data analysis

ATAC-seq was conducted as previously described.^54^ Briefly, 50,000 Nthy cells were collected and washed with ice-cold PBS. The supernatant was removed, and the pellet was resuspended in 50 µL of lysis buffer (10 mM Tris-HCl, pH 7.4, 3 mM MgCl_2_, 10 mM NaCl, and 0.1% NP-40). The nuclei pellet was then resuspended in 50 µL of transposition reaction buffer (10 mM TAPS, pH 8.5, 5 mM MgCl_2_, and 10% DMF) containing adaptor-loaded Tn5 enzyme (Enzynomics, EZ036) and incubated at 37°C for 30 minutes in a thermomixer set to 1000 rpm. After incubation, 10 µL of tagmentation clean-up buffer (900 mM NaCl and 300 mM EDTA, pH 8.0) was added to terminate the reaction. The tagmented DNA was purified using the MiniElute PCR Purification Kit (Qiagen, 28004) and quantified via qPCR. Library preparation was completed using the KAPA HiFi HotStart PCR Kit (Roche, KK2501) along with index primers^55^ and further purified with AMPure XP beads (Beckman, A63881). Libraries were sequenced in a 100 bp paired-end mode, with an Illumina NovaSeqX platform. The adapter sequences were trimmed using fastp,^56^ and reads were aligned to the hg38 and k12 reference genomes using Bowtie2.^57^ After alignment, mitochondrial reads and reads mapping to the ENCODE black list^58^ were removed. For downstream analysis, BAM files were normalized using RPM to account for differences in sequencing depth. Additionally, *E. coli* spike-in reads were used for further normalization to correct for technical variability. Bigwig files were generated using deepTools^59^ and visualized in IGV^60^. Heatmaps were also generated with deepTools to compare chromatin accessibility across conditions.

### CUT&Tag and data analysis

CUT&Tag was carried out following the Epicypher protocol. In brief, 50,000 Nthy nuclei and 5,000 *Drosophila* nuclei were incubated with ConA-coated magnetic beads (Epicypher, 21-1401) in bead activation buffer (20 mM HEPES-NaCl, pH 7.9, 10 mM KCl, 1 mM CaCl_2_, and 1 mM MnCl_2_) for 10 minutes at room temperature. The nuclei-bound ConA beads were then incubated with 0.5 µg of either H3K27me3 (CST, 9733) or H3K36me2 (ThermoFisher, MA-514876) and *Drosophila* H2Av antibody (Active Motif, 53168) diluted in antibody buffer (20 mM HEPES-NaCl, pH 7.9, 150 mM NaCl, 0.5 mM spermidine, 0.01% digitonin, 2 mM EDTA, and protease inhibitors), followed by overnight incubation at 4°C on a nutator (30 rpm). The next day, the complexes were washed twice using digitonin 150 buffer (20 mM HEPES-NaCl, pH 7.9, 150 mM NaCl, 0.5 mM Spermidine, 0.01% Digitonin, and protease inhibitors) and incubated with 0.5 µg of the secondary antibody diluted in digitonin 150 buffer for 30 minutes at room temperature. After washing twice with digitonin 150 buffer, 2.5 µL pAG-Tn5 (Epicypher, 15-1017) diluted in digitonin 300 buffer (20 mM HEPES-NaCl, pH 7.9, 300 mM NaCl, 0.5 mM spermidine, 0.01% digitonin, and protease inhibitors) was added, and the mixture was incubated for 1 hour at room temperature. After binding, the complexes were washed twice using digitonin 300 buffer, then incubated with digitonin 300 buffer supplemented with 10 mM MgCl_2_ for 1 hour at 37°C to tagment the target DNA. Following tagmentation, the complexes were washed with TAPS buffer (10 mM TAPS, pH 8.5 and 0.2 mM EDTA). SDS release buffer (10 mM TAPS, pH 8.5 and 0.1% SDS) was added, and the mixture was incubated at 58°C for 1 hour. After incubation, 0.67% triton X-100 was added, and the supernatant was transferred to a new PCR tube for non-hot start PCR, using 2x PCR master mix (Epicypher, 15-1018) with i5 and barcoded i7 primers. Subsequently, library DNA was purified with AMPure XP beads (Beckman, A63881). Libraries were sequenced in a 100 bp paired-end mode, with an Illumina NovaSeq 6000 platform. The adapter sequences were trimmed using Cutadapt,^61^ and reads were aligned to the hg38 and dm6 reference genomes using Bowtie2. After alignment, potential PCR duplication reads were removed using Picard. For downstream analysis, BAM files were normalized using CPM to account for differences in sequencing depth. Additionally, *Drosophila* reads were used for further normalization to correct for technical variability. H3K27me3 and H3K36me2 peak calling were performed using SICER2^62^ with a window size of 1,000 bp and a gap size of 100,000 bp. Bigwig files and heatmaps were generated with deepTools.

### ChIP-seq and data analysis

5×10^6^ Nthy cells were collected and crosslinked with 2 mM DSG (Thermo Fisher, 20593) for 35 minutes, followed by 1% formaldehyde for an additional 10 minutes. Crosslinking was quenched with 125 mM glycine, and the cell pellet was washed with PBS. After fixation, nuclei were isolated using a buffer containing 20 mM HEPES-NaCl, pH 7.9, 10 mM KCl, 0.1% triton X-100, 20% glycerol, 0.5 mM spermidine, and protease inhibitors. The nuclei were then resuspended in 130 μL of 1% SDS lysis buffer (50 mM Tris-HCl, pH 8.0, 10 mM EDTA, 1% SDS, and protease inhibitors) and sonicated using a Covaris S220 sonicator (175 peak incident power, 10% duty factor, 200 cycles per burst) for 430 seconds. The sheared chromatin was diluted in ChIP dilution buffer (16.7 mM Tris-HCl, pH 8.0, 0.01% SDS, 1.1% Triton X-100, 16.7 mM NaCl, 1.2 mM EDTA, and protease inhibitors) and centrifuged at 13,000 rpm for 15 minutes at 4°C to obtain the soluble chromatin fraction. Chromatin was incubated overnight at 4°C with Dynabeads Protein A (Thermo Fisher, 10001D) conjugated to EZH1 antibody (Proteintech, 20852-1-AP). The following day, the Dynabeads were washed three times with wash buffer A (140 mM NaCl, 1 mM EDTA, 0.5 mM EGTA, 1% triton X-100, 0.1% SDS, 0.1% sodium deoxycholate, and 10 mM Tris-HCl, pH 8.0), twice with wash buffer B (300 mM NaCl, 1 mM EDTA, 0.5 mM EGTA, 1% triton X-100, 0.2% SDS, 0.1% sodium deoxycholate, and 10 mM Tris-HCl, pH 8.0), and once with wash buffer C (250 mM LiCl, 10 mM Tris-HCl, pH 8.0, 1 mM EDTA, 0.5% NP-40, and 0.5% sodium deoxycholate). The beads were then resuspended in 10 mM Tris-HCl, pH 8.0 and incubated with 5 μg RNase A (Thermo Fisher, EN0531) for 30 minutes at 37°C, followed by incubation with 100 μg proteinase K (Thermo Fisher, EO0491) overnight at 65°C. The next day, the supernatant was transferred to new tubes, and size selection was performed using AMPure XP beads (Beckman, A63881) to purify DNA fragments around 200 bp in length. Library preparation was conducted using the NEB DNA library prep kit (NEB, E7645L). Libraries were sequenced in a 100 bp paired-end mode, with an Illumina NovaSeq 6000 platform. The adapter sequences were trimmed using Cutadapt, and reads were aligned to the hg38 reference genome using Bowtie2. EZH1 peak calling was performed using MACS2^63^ in narrow peak mode with a q-value threshold of 0.05. For additional downstream analysis, BAM files were normalized using CPM to account for differences in sequencing depth. Bigwig files and heatmaps were generated with deepTools.

### RNA-seq and data analysis

RNA was extracted as described in RT-qPCR method, and library preparation was performed using TruSeq Stranded Total RNA Ribo-Zero H/M/R Gold kit (Illumina, RS-122-2301). Sequencing was conducted in 100 bp paired-end mode on the Illumina NovaSeq 6000 platform. Adapter sequences were trimmed using Trimmomatic^64^, and reads were aligned to the hg38 reference genome using HISAT2.^65^ Transcript assembly was performed with StringTie^66^ for downstream analysis. Differential gene expression analysis was carried out using DESeq2 with *p*-value < 0.05 and log_2_ |fold change| > 1.^67^ For public RNA-seq data analysis, TPM counts for 513 thyroid tumor tissues and 686 thyroid normal tissues were obtained from TCGA and GTEx, respectively.

### Statistical analysis

Statistical significance was calculated using R and Prism software. Statistical differences were assessed using two-tailed Welch’s t-test or a two-tailed Student’s t-test. Additionally, one-tailed paired t-test was performed in Figure 5G. For all figures, asterisks denote statistical significance (\**p*<0.05. \*\**p*<0.01, \*\*\**p*<0.001, and \*\*\*\**p*<0.0001).

## Supplementary figures and figure legends

**Figure S1. The EZH1^Q571R^ mutation, commonly found in thyroid cancer, enhances H3K27 methylation level, related to Figure 1**.

**(A)** Western blot of β-tubulin, LMNA/C, H3, H3K9me3, and H3K27me3 in subcellular fractions of Nthy cell lines overexpressing EZH1^WT^ or EZH1^Q571R^.

**(B)** Bar graph representing the ratio of H3K27me3 (black), H3K9me3 (red), and H4K20me3 (blue) peptides in Nthy cells expressing EZH1^WT^ or EZH1^Q571R^. Data are presented as mean ± SEM, n=3 independent experiments. Statistical differences were assessed using two-tailed Welch’s t-test (\*\*\*\**p*<0.0001 and ns: not significant).

**Figure S2. Cells with EZH1^Q571R^ show stronger chromatin binding increased H3K27me3, chromatin condensation, and tumor suppressor gene suppression, related to Figure 2**.

**(A-D)** Principal component analysis plots showing the variance in H3K27me3 CUT&Tag (A), EZH1 ChIP-seq (B), ATAC-seq (C), and RNA-seq (D) across Nthy cells expressing EZH1^WT^ or _EZH1Q571R._

**(E)** Violin plots showing RNA expression of 24 genes in 686 thyroid normal tissues (green) and 513 thyroid tumor tissues (purple).

**(F)** Box plot showing RNA expression of the 119 EZH1^Q571R^-influenced downregulated genes and 5,369 genes in EZH1-bound regions in Nthy cells expressing EZH1^WT^ or EZH1^Q571R^.

**(G)** Top seven KEGG pathways enriched in the 5,369 genes.

**(H)** Representative Integrative Genomes Viewer tracks showing signals for EZH1 (red), H3K27me3 (blue), and RNA-seq (green) in Nthy cell lines expressing EZH1^WT^ or EZH1^Q571R^. The selected regions highlight known tumor suppressor genes that exhibit inherently low transcription under both conditions.

**Figure S3. PRC2-EZH1^Q571R^ significantly stimulates histone methyltransferase activity via enhanced nucleosome binding, related to Figure 3**.

**(A)** Histone methyltransferase (HMT) assays. Data are presented as mean ± SEM, n=3 independent experiments. Top: HMT assays were conducted in the presence of 300 ng mono-nucleosome and varying concentrations of PRC2 (160, 320, or 640 nM). H3K27me3 level was measured by western blot (top image) and the relative concentrations of PRC2 components (middle image) or mono-nucleosomes by Coomassie blue staining of SDS-PAGE gels (bottom image). Bottom: Quantification of the relative level of H3K27me3-modified mono-nucleosomes.

**(B)** Histone methyltransferase (HMT) assays. Data are presented as mean ± SEM, n=3 independent experiments. Top: HMT assays were conducted in the presence of 300 nM histone octamer and varying concentrations of PRC2 (200, 400, or 800 nM). Incorporation of [^3^H]-SAM into H3 was determined by autoradiography (top image) and the relative amount of PRC2 components by Coomassie blue staining of SDS-PAGE gels (bottom image). Bottom: Quantification of the relative level of [^3^H]-SAM-incorporated H3.

**Figure S4. PRC2-EZH1^Q571R^ compacts DNA much more efficiently than the wild-type, related to Figure 4**

**(A-B)** Fluctuation analysis and representative images. (A) Left, DNA images of the *i*-1^th^ frame (red), consecutive *i*^th^ frame (blue), and overlay merging those frames (white). (B) Depiction of the regions with variance between the *i-*1^th^ and *i*^th^ frames.

**(C-F)** Representative snapshots, kymographs, and real-time DNA fluctuation graphs of the EZH1 wild-type and various mutants.

**(G)** Density plot showing the frequency density of PRC2-EZH1^WT^, -EZH1^Q571R^, -EZH1^Q571A^, and - EZH1^Q571E^ over time.

**Figure S5. PRC2-EZH1^Q571R^ alters actively transcribed regions marked by H3K36me2/3, related to Figure 5**

**(A)** Histone methyltransferase (HMT) assays. Data are presented as mean ± SEM, n=3 independent experiments. Top: HMT assay was conducted in the presence of 1 ug either unmodified or H3K36me3-modified mono-nucleosomes and varying concentrations of PRC2 (200, 400, or 800 nM). Incorporation of [^3^H]-SAM into histone H3 or EZH1 was determined by autoradiography (top two images), and relative amounts of PRC2 components (middle image) and mono-nucleosomes (bottom image) by Coomassie blue staining of SDS-PAGE gels. Bottom: Quantification of the relative level of [^3^H]-SAM incorporated into unmodified or H3K36me3-modified mono-nucleosomes.

**(B)** Representative Integrative Genomes Viewer track showing H3K36me2, H3K27me3 peak regions and their intersections in Nthy cells expressing EZH1^WT^ or EZH1^Q571R^.

**(C)** Close-up cryo-EM structure showing the position of the EZH1 Q571 residue and H3K36 (left) or H3K36me3 (right). Cryo-EM structures are modified from 6WKR and 7KSR (left) and from 8VOB and 7KSR (right).

---

**Table S1.**

| Patient | Sex | Age at diagnosis | Pathology | Tumor size | Capsular invasion | Paired normal tissue origin |
| --- | --- | --- | --- | --- | --- | --- |
| #1 | Male | 44 | Minimally invasive follicular carcinoma | 4.2 x 3.5 x 3.0 cm | Present, minimal | Right thyroid lobe |
| #9 | Male | 54 | Minimally invasive follicular carcinoma, Oncocytic variant | 4.1 x 3.1 x 2.2 cm | Present, minimal | Left thyroid lobe |
| #52 | Female | 78 | Oncocytic adenoma | 3.5 x 2.2 x 2.0 cm | Absent | Total thyroid lobe |
| #53 | Female | 68 | Oncocytic carcinoma, minimally invasive | 5.6 x 2.8 x 2.3 cm | Present, complete | Right thyroid lobe |

