## Supplementary figures and images for "EZH1 Q571R-mediated chromatin compaction and its oncogenic potential in thyroid cancer"

### Supplementary figures

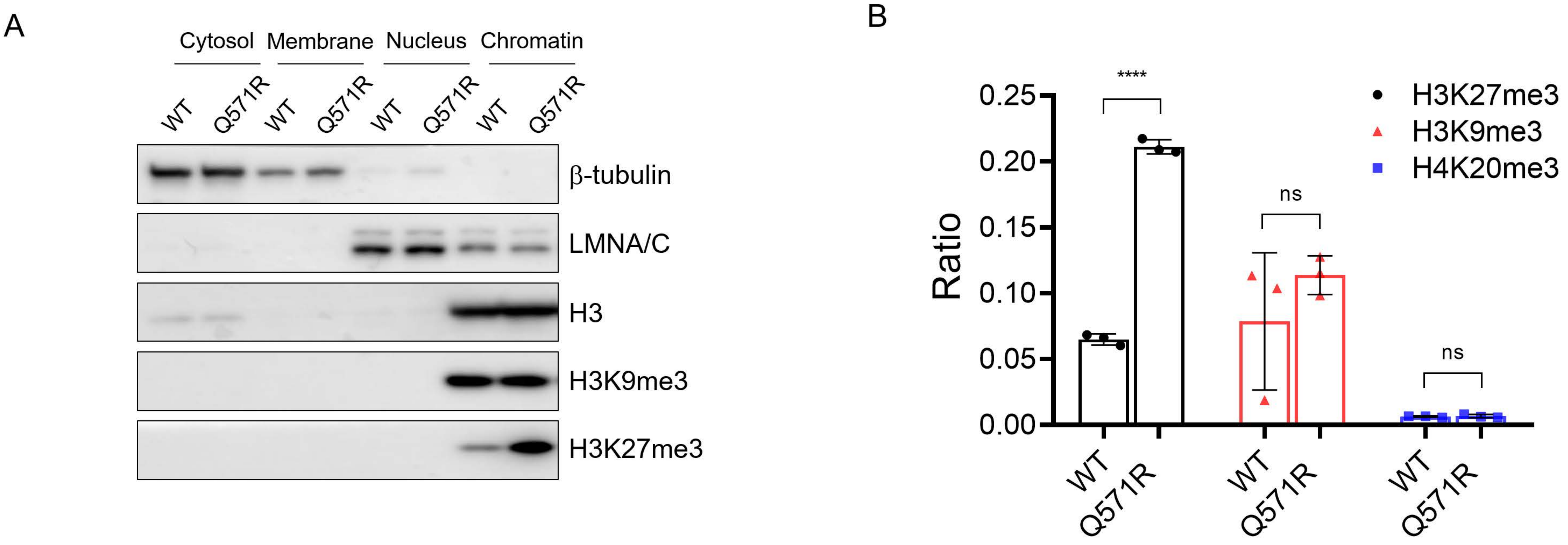

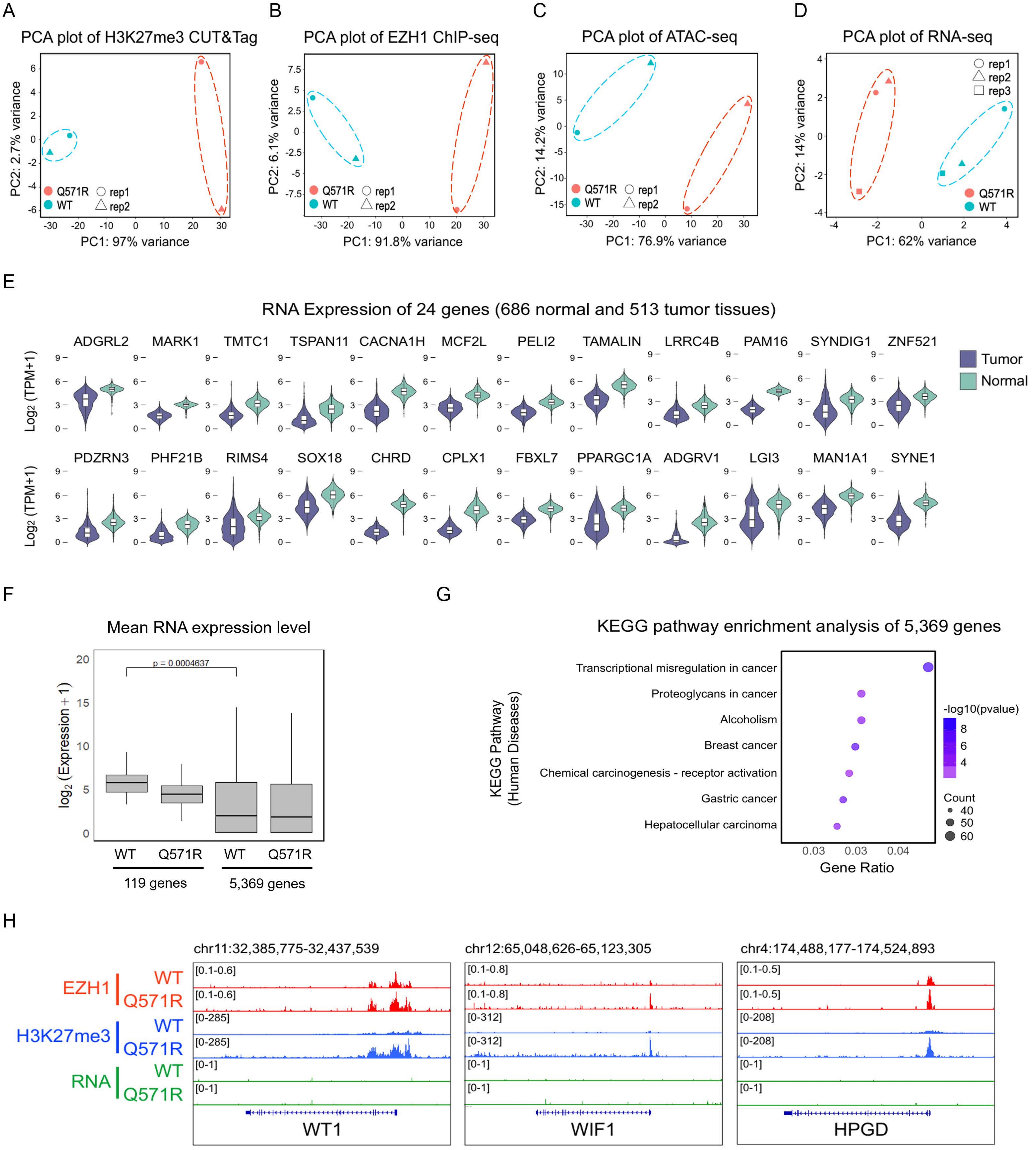

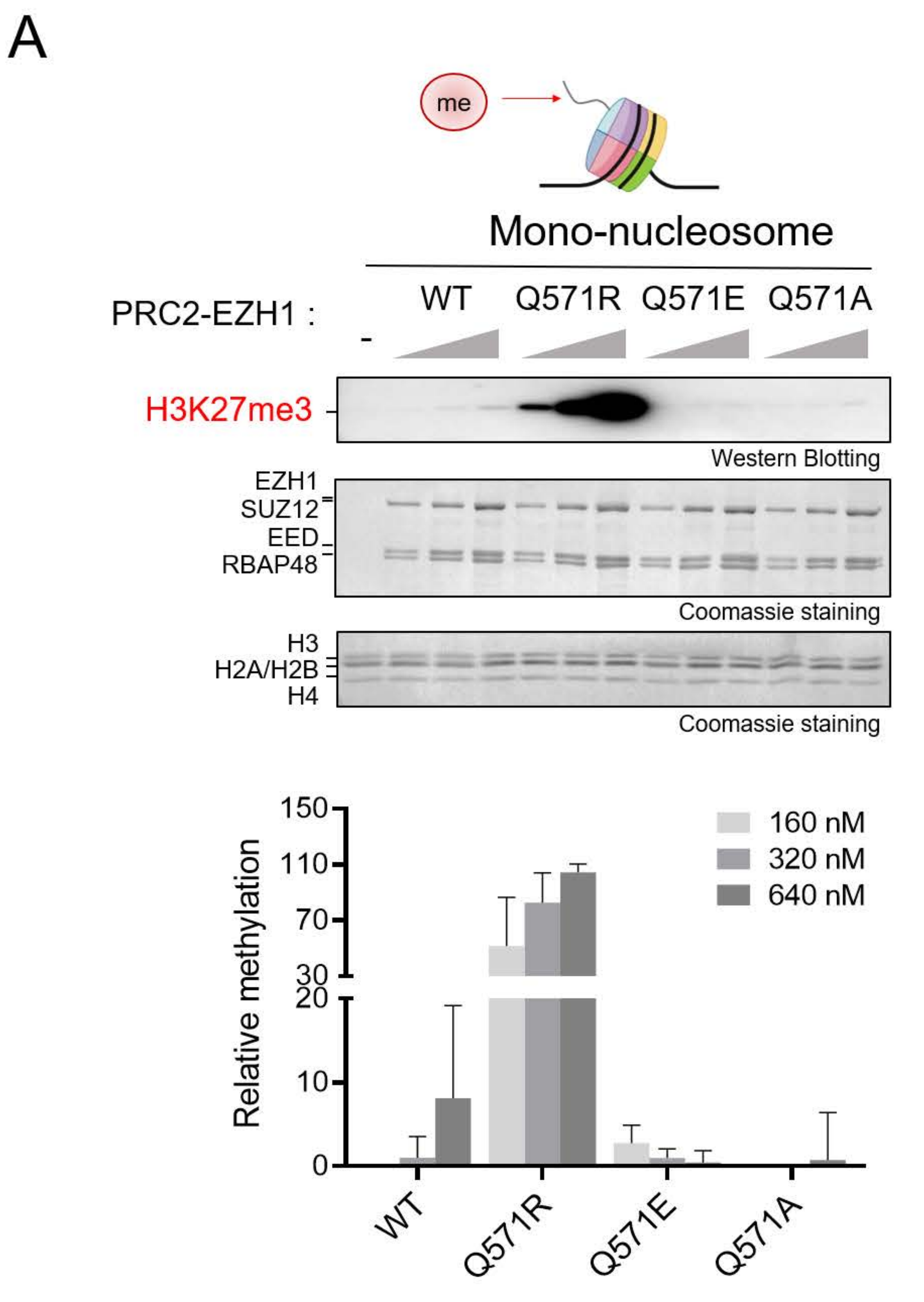

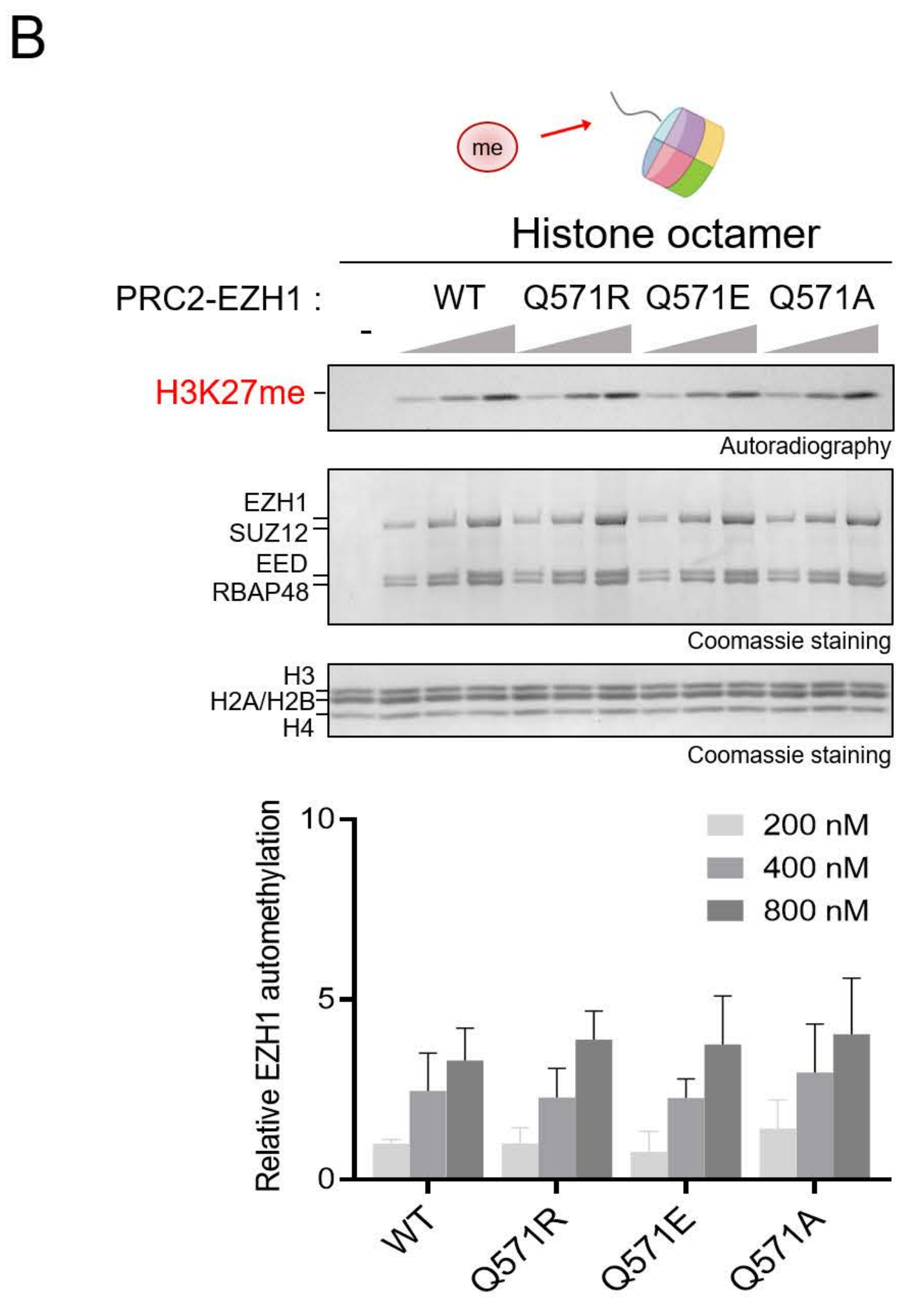

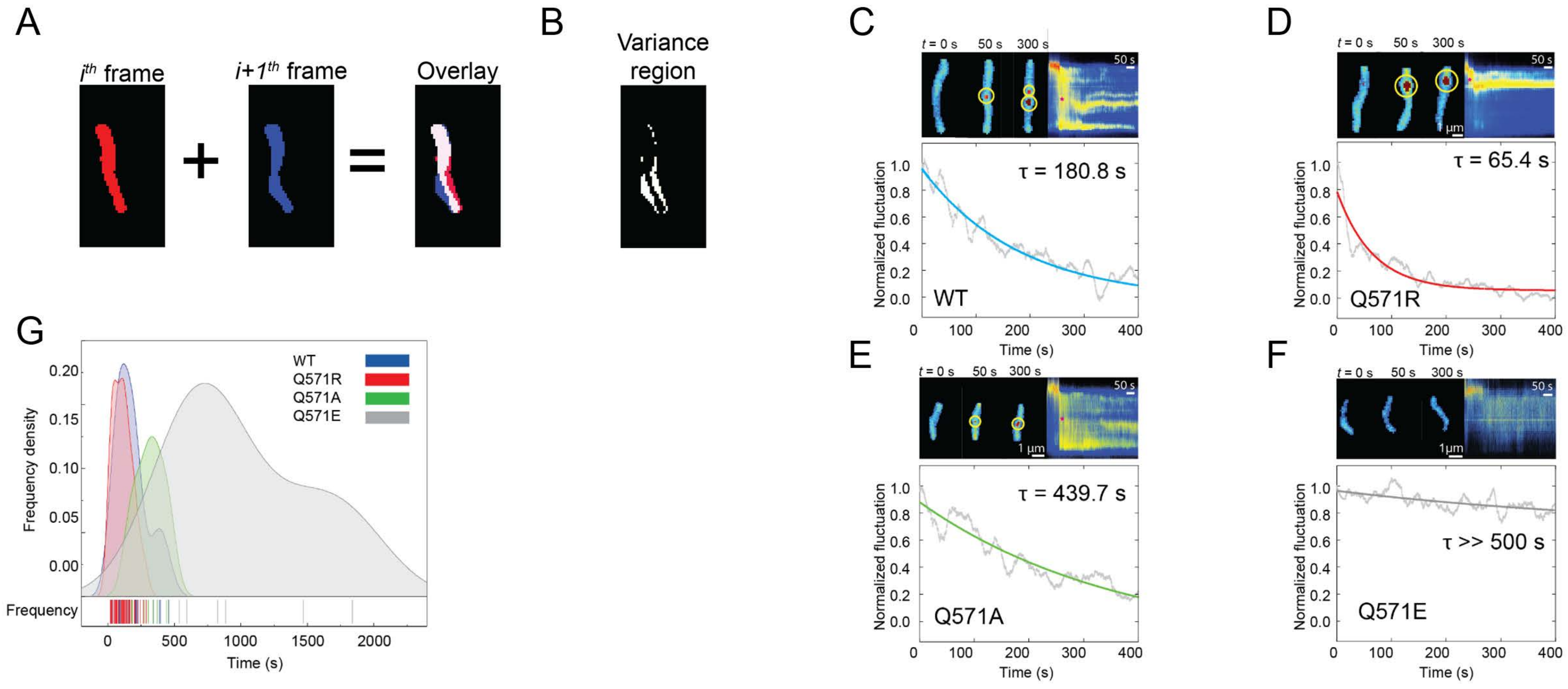

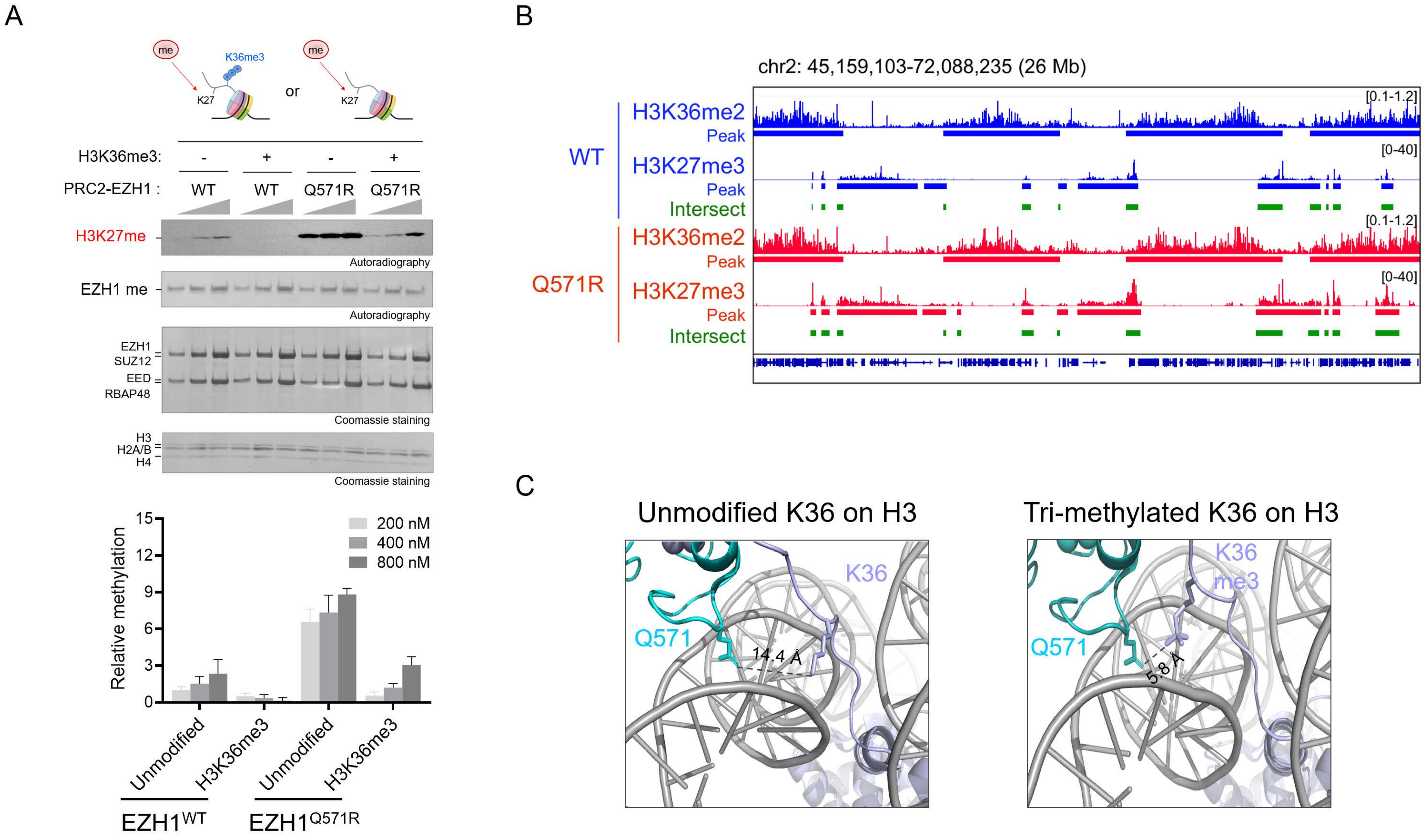
